# Single Cell Deconstruction of Muscle Stem Cell Heterogeneity During Aging Reveals Sensitivity to the Neuromuscular Junction

**DOI:** 10.1101/2020.05.28.121426

**Authors:** Peter J. Ulintz, Jacqueline Larouche, Mahir Mohiuddin, Jesus Castor Macias, Sarah J. Kurpiers, Wenxuan Liu, Jeongmoon J. Choi, Lemuel A. Brown, James F. Markworth, Kanishka de Silva, Benjamin D. Levi, Sofia D. Merajver, Joe V. Chakkalakal, Young C. Jang, Susan V. Brooks, Carlos A. Aguilar

## Abstract

During aging and neuromuscular diseases, there is a progressive loss of skeletal muscle volume and function in that impacts mobility and quality of life. Muscle loss is often associated with denervation and a loss of resident muscle stem cells (satellite cells or MuSCs), but the relationship between MuSCs and neural control has not been established. Herein, using a combination of single-cell transcriptomic analysis, high-resolution immunofluorescence imaging and transgenic young and aged mice as well as from mice with neuromuscular degeneration (Sod1^-/-^), a compensatory neuro-responsive function for a subset of MuSCs was identified. Genetic rescue of motor neurons in Sod1^-/-^ mice reduced this subset of MuSCs and restored integrity of the neuromuscular junction (NMJ) in a manner akin to young muscle. Administration of severe neuromuscular trauma induced young MuSCs to specifically engraft in a position proximal to the NMJ but in aging, this behavior was abolished. Contrasting the expression programs of young and aged MuSCs after muscle injury at the single cell level, we observed distinctive gene expression programs between responses to neuro-muscular degeneration and muscle trauma. Collectively, these data reveal MuSCs sense synaptic perturbations during aging and neuro-muscular deterioration, and can exert support for the NMJ, particularly in young muscle.

**Highlights:**

- Transcriptional landscapes of single satellite cells from different ages before and after injury as well as neurodegenerative models before and after nervous rescue
- A population of satellite cells reside in close proximity to neuromuscular synapse, which are lost with age
- Denervation promotes satellite cell engraftment into post-synaptic regions of young as opposed to aged muscle

## Introduction

Skeletal muscle atrophy and weakness are primary features of physical frailty, and are common among the elderly and patients afflicted with neuromuscular disorders^1^. The decline in the health and repair of skeletal muscle can be partially attributed to decreases in number and function of a population of resident stem cells called satellite cells^2^ or muscle stem cells (MuSCs). MuSCs respond to muscle damage via molecular changes^3^ that facilitate activation followed by differentiation. In aging however, MuSCs are lost, undergo exhaustion^4^ and inefficiently repair tissue after injury, further contributing to degeneration and sarcopenia^5,6^. The reduced capacity for MuSCs to function in the context of age-associated impairments^7^ due to intrinsic molecular changes remains unclear, owing to several factors such as heterogeneity^8,9^ of the MuSC pool. Cell-to-cell variation has been suggested to play a pivotal role in the aging process^10^ and influence clonal dynamics^11^, but the diversity of the MuSC compartment that forms during healing and how this heterogeneity varies across age has not been profiled. Single-cell expression profiling^12^ offers a powerful means to explore these actions as well as providing insights into how ensembles of cells act individually and/or together before and after injury^13^ to maintain healthy muscle into old age.

Tissue resident stem cells such as MuSCs critically depend on crosstalk with niche components^14^. The MuSC niche includes the multinucleated myofiber, vasculature^15,16^, and connective tissue cells, which can all be regulated by motor neuron (MN) integrity^17,18,19^. During aging, fundamental processes affecting resident non-myogenic cells are altered and drive pathological dysfunction. A well-documented example is degeneration of MNs and the neuro-muscular junction (NMJ), the synapse between MNs and myofibers. Perturbations to the NMJ has been shown to result in disruption of MuSC quiescence, and genetic ablation of MuSCs impairs the regeneration of the NMJ^15,20^ after nervous insult. These results suggest coordinative behavior between the NMJ and MuSCs but, how neural control influences MuSCs and aging remains an open question. Insight into the quantitative signaling networks that occur between cell types critical to the maintenance and regeneration of the nerve-muscle interface will significantly advance our understanding of cellular interactions and communication networks during aging and neuromuscular disease, and inform therapeutic strategies.

Herein, to further assess the connection between MuSCs and NMJ disruption that develops with age, we first contrast the gene expression programs of single MuSCs from mice of different ages. We observe that MuSCs from aged mice are less quiescent and a subset of MuSCs express markers suggestive of an interaction with the NMJ. We next show this subset of MuSCs is also present in a mouse model of neuromuscular degenerative disease (Cu/Zn superoxide dismutase deficiency – Sod1^-/-^). Genetic rescue of MNs in this model attenuated NMJ degeneration and decreased the presence of MuSC subsets expressing NMJ markers. We next demonstrate the specific contribution of MuSC derived progenitors at and in the vicinity of young NMJs after severe neuromuscular trauma. Last, we show the transcriptomes of single MuSCs responding during recovery from MN trauma are distinct from MuSCs exposed to muscle degeneration. Collectively, these data reveal MuSCs sense NMJ perturbations associated with aging, neuromuscular disease, and MN injury.

## Results

### Identification of an NMJ-Committed Subset of Muscle Stem Cells That Increases with Age

To understand how MuSCs change with age, MuSCs were extracted from lower hind limb muscles (tibialis anterior-TA and gastrocnemius-Gas) of young (2-3 months) and aged (22-24 mos) wildtype mice using fluorescent activated cell sorting^21^ (FACS, Fig. 1b), with both negative (Sca-1^-^, CD45^-^, Mac-1^-^, Ter-119^-^) and positive surface markers^22^ (CXCR4^+^ & β1-integrin^+^). To further verify these surface markers enriched a pure population of MuSCs, uninjured hindlimb muscles of transgenic mice harboring a Cre/LoxP-based system for MuSC lineage tracing Pax7Cre^ER/+^; Rosa26^mTmG/+^ (P7^mTmG^)^23^ were utilized. This system ubiquitously expresses a loxP-flanked membrane red reporter and after tamoxifen administration and excision, Pax7^+^ MuSCs and their progeny are tagged with a membrane-bound green fluorescent protein (mGFP). Both young and aged WT FACS-Sorted-MuSCs (FSMs) and P7^mTmG^ mononucleated cells from uninjured hindlimb muscles were subjected to droplet-based single-cell mRNA sequencing (scRNA-Seq)^24^ and a total of 23,065 single MuSCs were generated (3,725 young FSMs, 6,603 aged FSMs, and 12,737 P7^mTmG^ cells) yielding a mean of 6,150 unique molecular identifiers (UMIs) per cell after basic quality filtering. After filtering and non-linear dimension reduction through uniform manifold approximation and projection (UMAP) clustering^25^, nine clusters / cell types were observed (Supp. Figs. 1a-b). To further assess the purity of the FSMs, scRNA-Seq profiles of cells isolated from limb muscles across lifespan (3 mos - Tabula Muris^26^ and 24 mos - Tabula Muris Senis) were integrated and compared (Fig. 1c; 2,301 Tabula Muris and 11,895 Tabula Muris Senis cells, mean 4,418 UMIs per cell).

**Figure 1.**
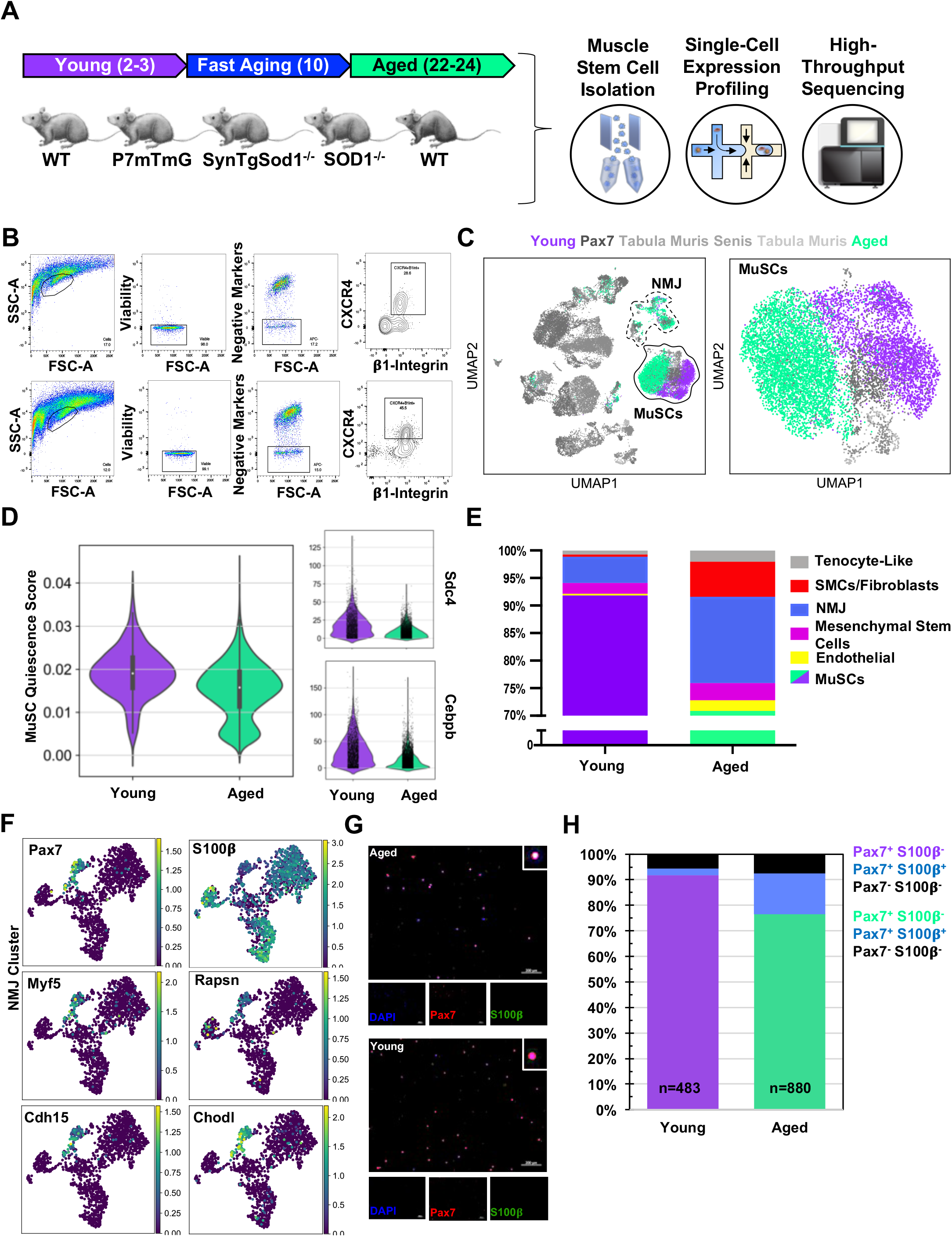
Skeletal muscle stem cells lose ability to retain quiescence in age and subsets of the cells adopt a neuro-regenerative response. A) Schematic of this study, whereby muscle stem cells (MuSCs) were FACS enriched from young (2-3 months) wild-type (WT) and Pax7^mTmG^ mice, 10-month old SynTgSod1 and SOD1 knockout mice, and aged (22-24 months) WT mice, then profiled using droplet-based single cell RNA sequencing (n=1 young and aged WT samples, n=2 Pax7^mTmG^ samples). B) Representative FACS scatter plots showing gating based on negative (Sca-1, Mac-1, CD45, Ter-119) and positive (CXCR4, B1-integrin) MuSC markers. C) Left-Dimensional reduction and unsupervised clustering of young WT FACS-isolated MuSCs, Pax7^mTmG^, aged WT FACS-isolated MuSCs, cells from limb muscle from Tabula Muris and Tabula Muris Senis datasets colored by sample type showing overlap between FACS enriched MuSCs and MuSC clusters among all mono-nucleated cells. Two independent replicates of each sample were sequenced where each sample is composed of a pool from two mice. Right-Re-clustering of MuSC cluster showing separation between young and aged MuSCs. D) Quiescence score calculated based on expression of 40 quiescence-associated genes suggest reduced quiescence among aged MuSCs (Mann–Whitney U = 3.7M, young = 2,725, aged = 4,606, p = 3.8e^−128^). (right) Normalized counts of Synedecan-4 (Sdc4) and CAAT/enhancer binding protein beta (C/EBPβ) showed reduced expression in aged MuSCs (p = 2.00e^−24^ and 1.63e^−60^, respectively). E) Stacked bar plot representing the fraction of the sorted cell populations that clustered with each mono-nucleated cell type. A higher fraction of FACS purified cells from aged mice are associated with the NMJ cluster as compared to young. F) Re-clustered UMAP diagrams of the NMJ-associated cluster and overlay of specific genes. G) Immunofluorescent stains against S100β and Pax7 among FACS enriched MuSCs. H) Quantification of S100β and Pax7 proteins show less than 5% of young MuSCs co-express both markers compared to 15% of aged MuSCs.

Integration of the different datasets revealed the transcriptomes of FSMs from different ages clustered and overlapped with unsorted MuSCs (Fig. 1c). Given the limited capacity of scRNA-Seq to enumerate differences in the activity of lowly expressed TFs (such as Pax7) that would contribute to differences in states such as maintenance of quiescence, we developed a quiescence score^27^ based on the expression of 40 quiescence-associated genes and contrasted young and aged FSMs. Young MuSCs displayed stronger quiescent scores when compared to aged MuSCs (Fig. 1d), which is consistent with previous studies showing aging perturbs the ability of MuSCs to retain a quiescent state^14^. Drivers of the stronger quiescent score in young MuSCs included genes that help restrain differentiation (Fig. 1d), such as syndecan 4, a heparin sulfate proteoglycan, and CAAT/enhancer binding protein beta (C/EBPβ)^28^, a transcription factor that inhibits MyoD. Comparison of the expression of different surface markers^3^ for the four different samples revealed highly similar profiles (Supp. Fig. 1c), and overlap between the samples and Tabula Muris and Tabula Muris Senis datasets suggests that FSMs capture the diversity of unsorted cells.

Quantification of the number of cells that clustered with MuSCs compared with other clusters revealed ~92% of young FSMs overlapped with the MuSC cluster (Fig. 1e). In contrast, ~73% of aged FSMs overlapped with the MuSC cluster, and annotation of the clusters based on their unique gene expression profiles revealed the largest non-MuSC cluster of FSMs expressed genes nominally associated with NMJ maintenance and repair such as S100β, a Ca^2+^ binding neurotrophic factor that induces neurite growth^29^ and is induced after peripheral nerve injury as well as Cadm1^30^, a homophilic cell adhesion molecule that promotes synapse formation^31^ and interacts with synaptic proteins^32^. Re-clustering of cells in the NMJ-associated cluster showed Pax7^+^ cells within this cluster also expressed Myf5, a myogenic transcription factor, M-cadherin (Cdh15), and chondrolectin (Chodl), a lectin that has been observed to mediate axonal growth and growth cone interactions^33^, as well as exhibit dysregulation in mouse spinal muscular atrophy^34^.

This subset of cells also expressed Rapsyn (Rapsn), an intracellular post-synaptically expressed protein that is essential for acetylcholine receptor (AChR) clustering and formation of NMJs (Fig. 1f) and has previously been observed to increase in aging and concentrate at the NMJ^35^. To rule out if FACS was enriching other cells types found at NMJs such as Schwann cells with MuSCs, young and aged FSMs were co-stained for Pax7 and S100β. Immunostaining of young and aged FSMs with Pax7 showed ~94% and ~93% were Pax7^+^, respectively (Fig. 1g-h). Young FSMs displayed a small fraction of Pax7^+^ / S100β^+^ cells (~2%), which was in contrast to the larger fraction of Pax7^+^ / S100β^+^ (~16%) observed in aged FSMs, and these results were approximately consistent with the percentages identified through scRNA-Seq. We also did not observe Pax7^+^ cells capping nerve terminals after denervation or in aging^17,18^, consistent with the known specificity of Pax7 for MuSCs in postnatal muscle^36^. Overall, these results confirm aging is associated with NMJ disruption and suggest that aging induces an increase in a subset of MuSCs that display commitment towards an NMJ phenotype.

### Motor Neuron Degeneration Induces NMJ-Associated Muscle Stem Cell Phenotypes

Since NMJ disruption is a feature of aging skeletal muscle, we asked if an orthogonal model of neuro-degeneration that triggers bouts of NMJ denervation and reinnervation with eventual failure would induce MuSC subsets with an NMJ committed phenotype. Sod1 is an enzyme that breaks down superoxide radicals and protects from oxidative damage, and Sod1^-/-^ mice (where Sod1 is globally knocked out) display NMJ degeneration and muscle loss early in life (Fig. 2a)^37^. Hindlimb muscles from 10-month old Sod1^-/-^ mice, which displayed denervation in even higher proportions than aged mice (data not shown), were harvested. MuSCs were isolated with FACS and profiled with scRNA-Seq recovering 6,575 cells. Similar to aged MuSCs, Sod1^-/-^ FSMs contained fractions of cells that did not cluster with other MuSCs (Fig. 2b), and the next largest Sod1^-/-^ cell cluster contained genes associated with NMJ function (Fig. 2c). IF imaging of Pax7 in Sod1^-/-^ FSMs revealed ~97% were Pax7^+^, consistent with the results from young and aged FSMs, ruling out contamination of other NMJ cellular components through FACS (Supp. Fig. 2). Computation of the quiescence score for Sod1^-/-^ FSMs revealed a similar score to aged FSMs (Fig. 3d). Both Sod1^-/-^ and aged MuSCs displayed stronger expression of markers associated with early activation (Itm2a, Supp. Fig. 2, increased in Sod1^-/-^ (mean=1.2) and aged MuSCs (mean=1.3) compared to young (mean=0.5)), and differentiation such as the members of a canonical Wnt signaling program (Dishevelled 1 - Dvl1, RhoA, and GSK3-β), as well as lower expression in adenomatous polyposis coli^38^ - APC, Csnk1a1 and Axin1 (Fig. 2e). Sod1^-/-^ and a small fraction of aged FSMs also contained a subset of progenitors that overexpressed Myogenin^39^ (MyoG), a developmental regulator that is upregulated in denervated muscle^40^, as well as Dok-7, a post-synaptic docking protein of MuSK^41^ (Fig. 2f). Integrating these data shows degeneration of MNs from Sod1^-/-^ results in modulation of MuSCs in a similar manner to aging, whereby MuSCs exhibit a pseudo-activation state and a subset of cells enact a program consistent with NMJ disruption.

**Figure 2.**
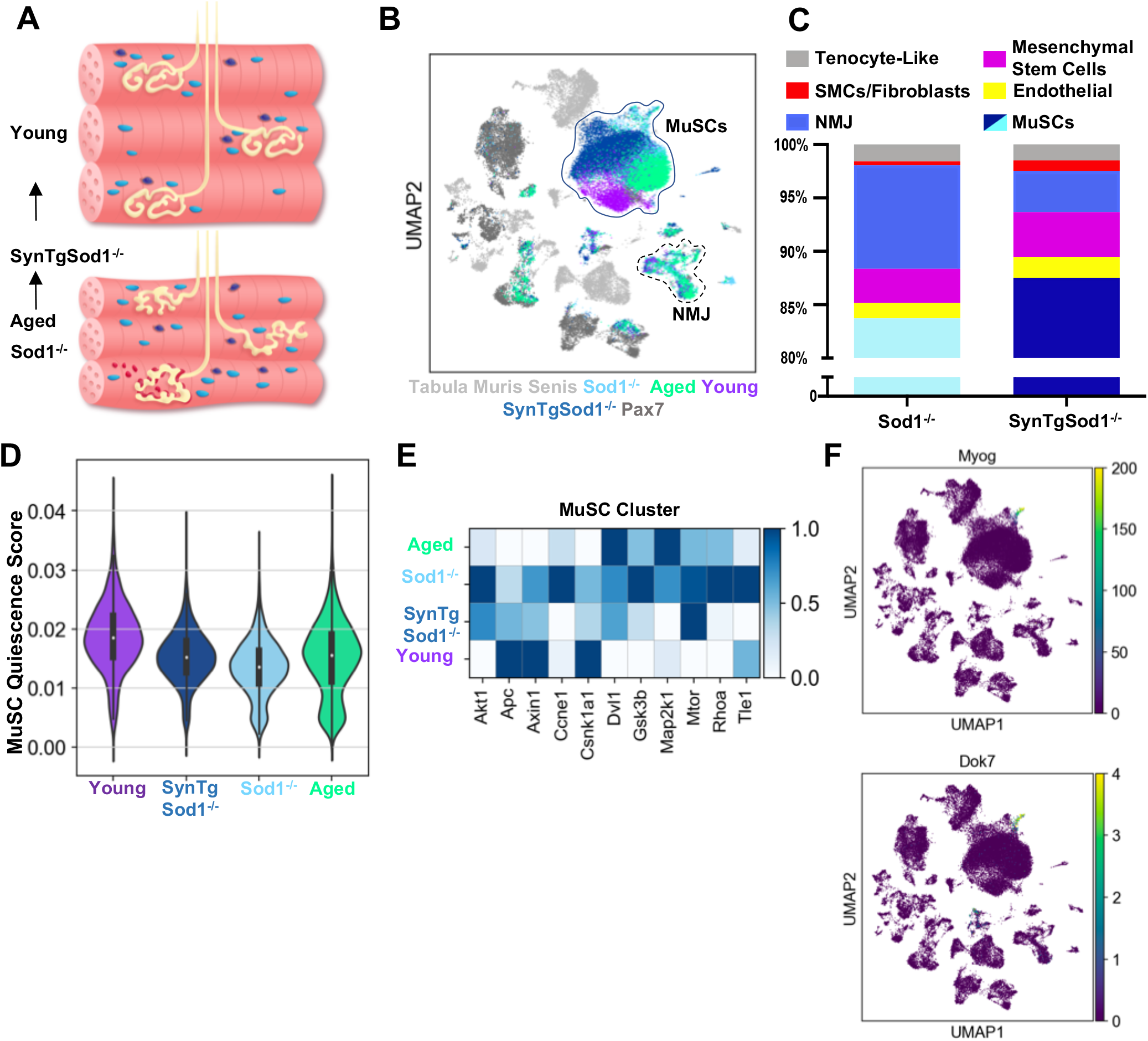
Neurodegeneration activates muscle stem cells in a similar manner to aging and rescue of motor neurons partially reverses this activation state. A) Schematic of neuromuscular junction (NMJ) in young (top) and aged or Sod1^-/-^ (bottom) muscle, whereby NMJ becomes fragmented or partially denervated. B) UMAP dimensional reduction of FACS-isolated MuSCs (FSMs) from young and aged WT mice, 10 month Sod1^-/-^ mice, 10 month Sod1^-/-^ rescue (SynTgSod1^-/-^) mice, Pax7^mTmG^ mice, and public Tabula Muris Senis limb muscle datasets colored by sample type. The main MuSC and NMJ clusters, identified via marker genes, are demarked by solid and dashed lines, respectively. Full unsupervised cluster designations for this UMAP diagram are provided in Supplemental Figure 3. C) Stacked bar plot representing the fraction of the sorted cell populations that clustered with each mono-nucleated cell type. More FACS purified MuSCs from Sod1^-/-^ mice cluster with NMJ-associated cells when compared to SynTgSod1^-/-^ mice. D) Quiescence score calculated based on expression of quiescence-associated genes show Sod1^-/-^ FSMs display similar quiescence mean score to aged MuSC; however, SynTgSod1^-/-^ also display reduced quiescence scores than young. E) Scaled expression (to a 0-1 scale) heatmap of MuSC cluster comparing Wnt signaling proteins from different samples displaying upregulation in Sod1^-/-^ and aged MuSCs compared to young and SynTgSod1^-/-^ MuSCs. F) Overlay of expression of Myogenin (Myog-top), a myogenic transcription factor that is upregulated in denervation and activates expression of synaptic proteins, and Docking protein 7 (Dok7-bottom), an adaptor protein of MuSK that promotes synaptic formation.

**Figure 3.**
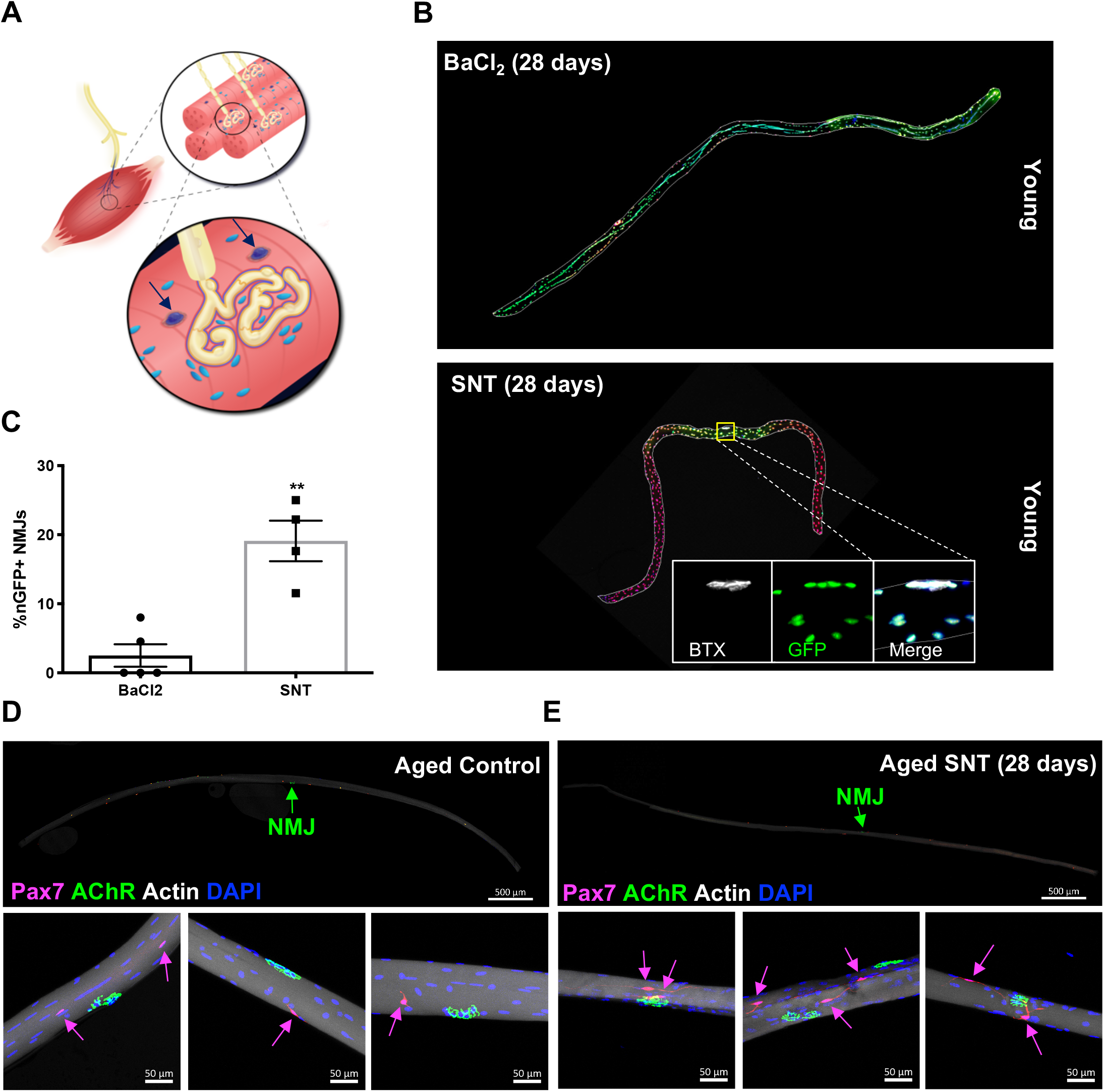
Denervation induces muscle stem cell actions proximal to the neuromuscular junction that is attenuated with age. A) Schematic of neuromuscular junction whereby muscle stem cells (MuSCs) labeled with dark blue arrows are in close proximity. E) Representative immuno-fluorescence images of single myofibers isolated from extensor digitorum longus (EDL) muscles from young Pax7^CreER/+^-Rosa26^NuclearTdTomato-NuclearGFP^ (P7^nTnG^) mice, which display red fluorescent protein (RFP) in their nuclei and following administration of tamoxifen, Pax7^+^ MuSCs and their progeny are labeled with a nuclear green fluorescent protein (GFP). After muscle injury through barium chloride (BaCl_2_) injection (top), myofiber degeneration resulted in contribution of MuSC derived progenitors and centrally located myonuclei along the entire length of the regenerated myofiber. After sciatic nerve transection (SNT), SNT MuSC derived myonuclei were confined at or near NMJ myofiber regions. Magnified inset images show NMJs (Btx, AChRs) with nGFP+ nuclei. Scale bar for myofibers = 200 μm for inset = 25 μm. n=5 for both injury types and 20-30 myofibers counted from each isolated muscle. C) Quantification of nGFP^+^ near synapses and in the mid regions of myofibers, where **p < 0.05 using two-sided t-tests. D) Representative immuno-fluorescence image of single tibialis anterior myofiber (top) from aged (20 mos, n=3) Pax7^CreER/+^-Rosa26^TdTomato/+^ mouse with neuromuscular junction (NMJ) labeled with green arrow. Pax7-Pink, Acetylcholine receptor (AChR) labeled with a-bungarotoxin (BTX) stain (white) and nuclei/DAPI (blue). Magnified images of NMJ from single myofibers (bottom) and MuSCs labeled with pink arrows. E) Representative immuno-fluorescence image of aged (20 mos, n=3) denervated single myofiber from Pax7^CreER/+^-Rosa26^TdTomato/+^ tibialis anterior muscle. Magnified images of NMJ from single myofibers (bottom) shows MuSCs in close proximity to the NMJ but do not engraft and develop long projections.

### Rescue of Degenerative Motor Neurons Partially Reverts Muscle Stem Cell Dysfunction

To determine if genetic rescue of MNs in Sod1^-/-^ mice would revert the MuSC response observed in Sod1^-/-^ and aged muscles, Sod1 was specifically overexpressed only in MNs via the synapsin 1 promoter (SynTgSod1^-/-^)^42^ and Sod1 was knocked out in other tissues. Previously, we showed this rescue prevents NMJ degeneration and muscle fiber denervation that is observed in Sod1^-/-^ mice^41^. Hindlimb muscles from age-matched SynTgSod1^-/-^ mice (10 mos) were harvested, MuSCs extracted with FACS and profiled with scRNA-Seq, recovering 10,939 cells. SynTgSod1^-/-^ FSMs revealed stronger overlap with young FSMs and a decrease in the percentage of FSMs that clustered with other cell types such as NMJ-associated cells (Fig. 2b-c). Enumeration of the quiescent score for SynTgSod1^-/-^ FSMs did not significantly vary when compared to aged and Sod1^-/-^ FSMs (Fig. 2d), but this effect may be due to the fact that Sod1 was lost in all other tissues including MuSCs. However, the expression of factors such as Zbtb20^43^ and Gpx3, a retinoic acid target that promotes stem cell self-renewal^44^, were increased in SynTgSod1^-/-^ FSMs compared to Sod1^-/-^ (Supp. Fig. 2d). SynTgSod1^-/-^ FSMs also exhibited a lack of subsets that expressed MyoG and Dok7, and reduced expression of canonical Wnt signaling genes (Fig. 2e-f). Amalgamating these results shows a subset of MuSCs sense and respond to the NMJ, and prevention of NMJ degeneration attenuates the activation of an NMJ-committed phenotype.

### Denervation Engenders Muscle Stem Cell Actions Proximal to NMJs

We reasoned that if MuSCs adopt an NMJ-committed state, acute perturbations to the NMJ would engender MuSC actions proximal to this niche (Fig. 3a). We contrasted acute and specific perturbation of MNs using sciatic nerve transection (SNT), with intramuscular injection of barium chloride (BaCl_2_). In the former, MuSCs demonstrated local roles in the reinnervation of young NMJs^18^ including contribution of new post-synaptic myonuclei; whereas the latter induces degeneration and MuSC-derived progenitor contribution along the length of the entire myofiber. We utilized a MuSC lineage tracing system (Pax7^CreER/+^-Rosa26^nTnG/+^: P7^nTnG^) whereby all nuclei contain a red fluorescent protein and after administration of tamoxifen, Pax7^+^ MuSCs and their progeny are indelibly labeled with a nuclear green fluorescent protein (nGFP)^17^. We have previously shown that after tamoxifen administration, only Pax7^+^ MuSCs are initially nGFP labelled along the length of isolated myofibers, and devoid of precocious nGFP expression in myonuclei^17^. Examination of single myofibers after both injuries (28 days) displayed variations in MuSC derived contribution of nGFP. As expected, BaCl_2_ induced myofiber degeneration results in contribution of MuSC derived progenitors and centrally located myonuclei along the entire length of the regenerated myofiber (Fig. 3b). In contrast, after SNT MuSC derived myonuclei were confined at or near young NMJ myofiber regions (Figs. 3b-c); consistent with previous reports whereby MuSC depletion leads to loss of myonculei in the vicinity of young NMJs after SNT^18^. Taken together, these results show MuSCs are sensitive to NMJ disruptions, and in such contexts, can give rise to progenitors that are presumably fated towards NMJ maintenance and repair given their local contribution in the vicinity of young NMJs.

To further evaluate the relationship of MuSCs with NMJ disruption and local changes in MuSC-derived contribution to myofibers in aging, we administered tamoxifen and performed SNT on aged (20 mos) Pax7^CreER/+^-Rosa26^TdTomato/+^ (P7^TdT^) tibialis anterior muscles. The TdTomato reporter is cytosolic; thus, if any MuSC derived cell fusion occurs diffuse reporter label will be observed in the myofiber. We extracted aged denervated and uninjured single muscle fibers after 28 days post injury, and observed fragmented and diffuse staining of post-synaptic nicotinic receptors, as well as smaller myofiber cross-sectional areas for aged SNT myofibers (data not shown). Uninjured MuSCs were observed along the length of the fiber with a small percentage (~6%) that were proximal to the NMJ (<50μm, Fig. 2d), which is consistent with previous studies^17,45^. In contrast, after SNT, aged MuSCs did not engraft into myofibers as evidenced by the lack of diffuse TdTomato reporter labeling. Rather, TdTomato was found restricted to aged MuSCs that were observed to migrate towards the NMJ and displayed elongated projections^46^ (Fig. 2e). These results suggest MuSCs are recruited to the NMJ as a result of MN perturbation, but restrained from differentiating and fusing into new synaptic myonuclei.

### Muscle Injury Induces a Continuum of Regenerative States in Muscle Stem Cells That is Distinct from Activation through Neural Degeneration

The variation in engraftment behavior of young and aged MuSCs has previously been observed after muscle injuries. To determine if acute muscle injury induced a subset of MuSCs to adopt an NMJ-committed state similar to that developed during aging and with chronic neuro-degeneration, hind limb muscles of mice were injured via BaCl_2_ injection. Histological and immunofluorescence (IF) imaging revealed destruction of the tissue (Fig. 4a) and regeneration. MuSCs were purified with FACS at multiple time points (3 days, 7 days post injury-dpi) and subjected to scRNA-Seq, which recovered 7,441 MuSCs after filtering. These datasets were combined with uninjured young FACS-sorted MuSCs (3,725 cells) and the Pax7^mTmG^ and Tabula Muris hindlimb public datasets were once again added. UMAP dimension reduction and clustering revealed separation of cells by day of isolation (Fig. 4b). These data show MuSC heterogeneity as a result of injury but distinct separation from the NMJ-associated cluster (Supp. Figure 4a). Quantitation of the number of young MuSC in each identified cluster during the regenerative process revealed a decrease in the percentage of FSMs that overlapped with the NMJ-associated cluster for 3 and 7 dpi, respectively (Supp. Fig. 4b).

**Figure 4.**
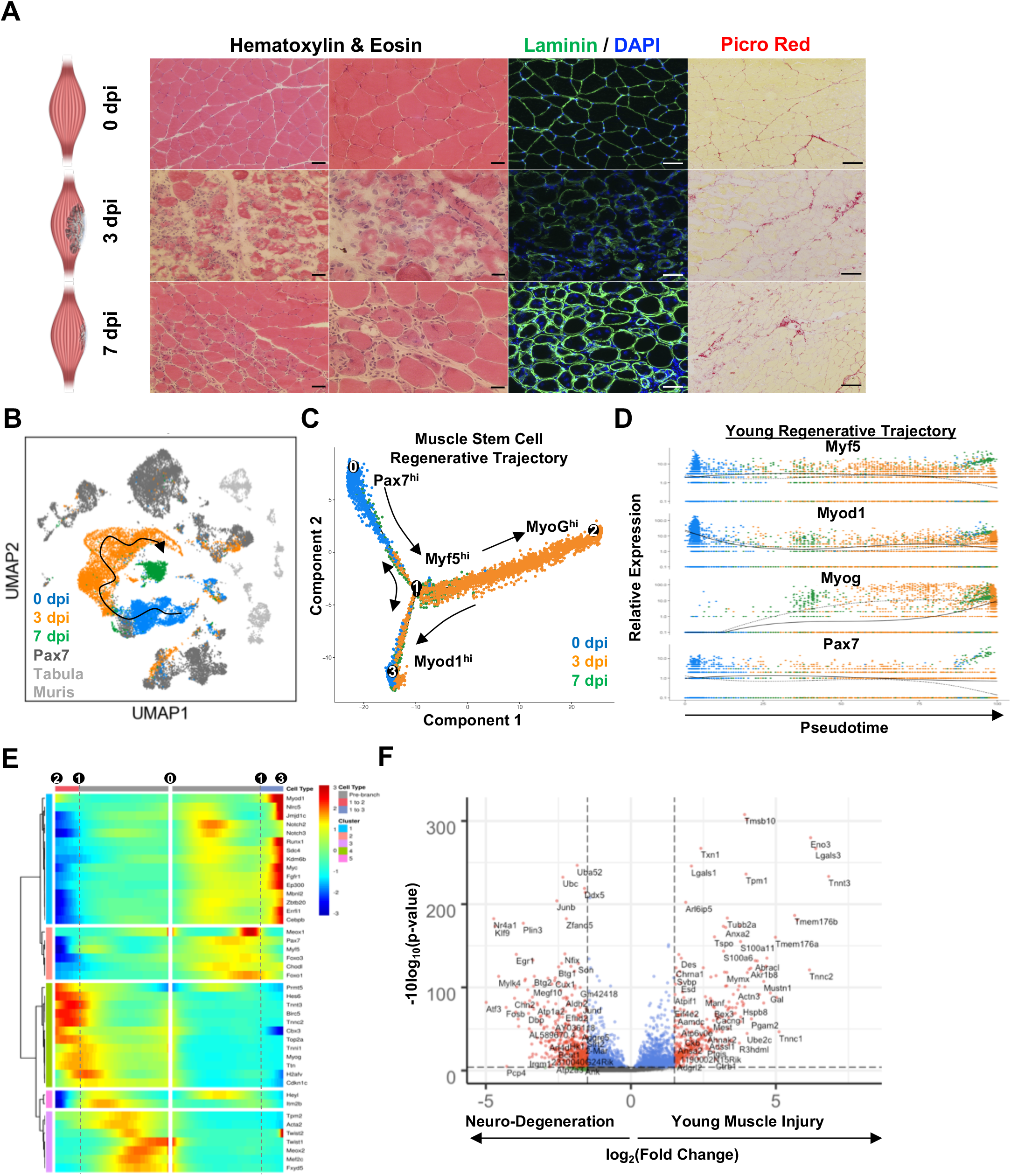
Single-cell transcriptomes of young muscle stem cells during muscle regeneration reveal heterogeneity but distinct regenerative dynamics compared to muscle stem cells isolated in response to neuro-degeneration. A) Histological cross-sections of tibialis anterior muscles isolated before and after injury (BaCl_2_) at 3 and 7 days post injury (dpi). Crosssections are stained with Hematoxylin and Eosin (H&E), Laminin and DAPI, Picrosirius Red bright field showing destruction of tissue and regeneration. B) UMAP dimensional reduction of young FACS-isolated MuSCs isolated before (0dpi-blue) and after injury (3dpi-orange, 7dpi-green) as well as unsorted Pax7^mTmG^ MuSCs (dark gray), cells from limb muscle from 3-month-old Tabula Muris datasets (Light Gray). A possible developmental trajectory suggested by expression of canonical myogenic factors plotted in c) is indicated by the arrow. C) Pseudo-time reconstruction of muscle stem cells before and after injury labeled by day of isolation (0 – blue, 3 dpi - orange, 7 dpi - green) displays a branched trajectory also reflective of quiescent, activated, differentiated and returning towards quiescence myogenic states. The relative peaks in expression of transcription factors Pax7 (state 0-1), Myf5 (state 0-1), Myod1 (state 1-3) and MyoG (state 1-2) are labeled on the trajectory. Arrows are labeled as guides. D) Expression tracking of myogenic regulatory factors Pax7, Myf5, MyoD1 and Myog during trajectory, where each dot represents a single cell color-coated by day of isolation (blue-0dpi, orange-3 dpi, green-7dpi). The black solid line corresponds to trajectory (0-1-2) on (C) and dashed line corresponds to trajectory (0-1-3). E) Split heatmap of genes differentially expressed in pseudotime, where the beginning of the pseudotime trajectory is in the middle of the diagram (point ‘0’), and the two branch trajectories extending left and right to endpoints ‘2’ and ‘3’, respectively. Branchpoint ‘1’ is indicated in both trajectories by vertical dotted lines. Rows in the heatmap are hierarchically clustered into five general groups based on expression. F) Volcano plot displaying differential expression of genes from MyoG^hi^ MuSCs from state 2 after muscle injury (293 cells) to MyoG^hi^ MuSCs isolated from neurodegeneration (Sod1^-/-^) and age (155 cells). Significantly differentially expressed genes highlight differences between a healthy myogenic repair program adopted by young MuSCs after muscle injury vs the activation program of Sod1^-/-^ and aged MuSCs.

To probe further into drivers of the variance between clusters during regeneration, MuSCs were ordered in pseudotime^47^ (Fig. 4c). Cells from 0 dpi clustered towards the root of the trajectory (states 0 to 1) with a branch that contained a subset of activated cells (state 3). In contrast, cells from 3 dpi occupied the end of the continuum (state 2). Activated cells from 7 dpi spanned an intermediate space indicating a return towards the expression state of cells sampled from 0 dpi. Overlay of canonical myogenic regulatory factors (Pax7, Myf5, Myod1, MyoG) onto the trajectory demonstrated MuSCs in different states (Fig. 4d). For example, Pax7^+^ MuSCs were enriched at 0 and 3 dpi and MyoG^+^ MuSCs were concentrated in 3 dpi samples. Cells from each day were heterogenous, with cells from 0 dpi containing both quiescent (state 0: Pax7^hi^ / MyoD^low^) and activated (state 3: Pax7^low^ / MyoD^hi^) cells. These results were consistent with upregulation of factors that are well known components of the quiescent state (Fig. 5e, Supp. Fig. 4c), such as Sdc4, Chodl, Fgfr1, C/EBPβ, Foxo1 and Foxo3 (TFs that activate antioxidant genes and protect the quiescent state^48^). As cells move away from the root at 3 dpi (transitioning from states 1 to 2 on Fig. 5d), genes associated with quiescence were downregulated and increases in expression of genes nominally associated with activation were observed such as Myf5, c-Myc, HeyL, Ep300, Runx1, H2afv, Birc5, Top2a, and Nlrc5 (Fig. 4e), which has been shown to be upregulated in proliferating MuSCs and downregulated in quiescent MuSCs that evade immune surveillance^49^. Further along the trajectory, early myogenic commitment factors such as Mef2c, Itm2b, Acta2, Tpm2, and Fxyd5, a negative regulator of cell adhesion, peaked in expression. Towards the end of the trajectory, markers associated with differentiation, such as Hes6, Tmem8c, and Cdkn1c (p57) and as well as sarcomeric proteins (Acta2, Ttn, Myh3, Tnni1, Tnnt3, Tnnc2) were enriched. MuSCs from 7 dpi spanned an intermediate space between the start and end of the trajectory and increased expression of collagen V factors, which have been shown to influence the quiescent phenotype^50^, splicing regulators such as muscle-blind like protein 2 (Mbnl_2_), several demethylases (Jmjd3/Kdm6b and Jmjd1c) that remove methyl marks from histone 3 on lysine 27 (H3K27) and Prmt5 (a regulator of MuSC proliferation^51^). These cells also upregulated Notch genes (Notch2, Notch3), Sprouty 2 (Spry2), and Zbtb20. Integrating these data shows MuSCs exhibit heterogeneity during regeneration and utilize multiple factors to execute stage-specific transitions such as differentiation.

**Figure 5.**
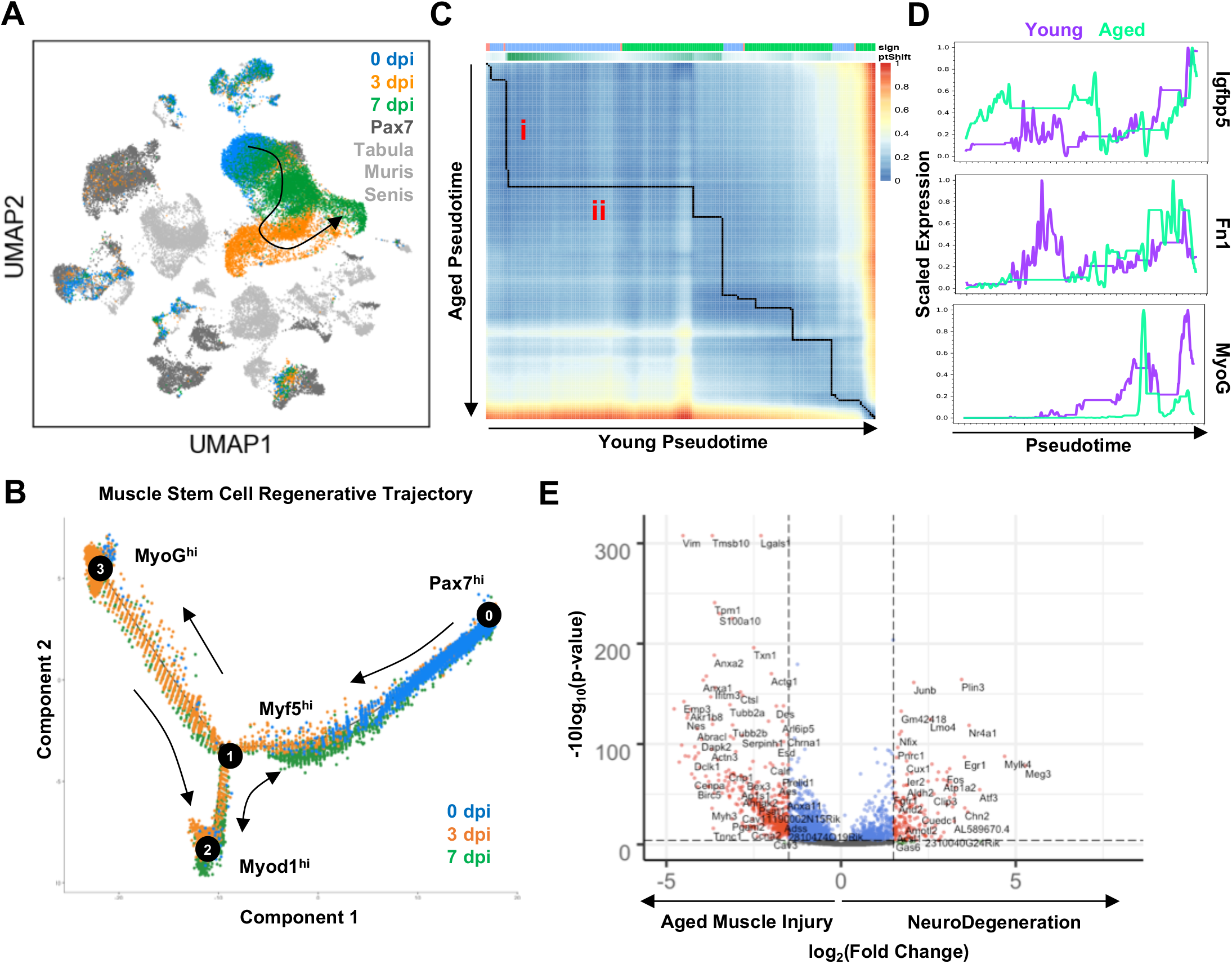
Expression profiling of single muscle stem cells from aged mice during muscle healing shows regenerative defects and distinct gene expression programs compared to muscle stem cells responding to neuro-regeneration. A) UMAP dimensional reduction of aged FACS-isolated MuSCs at timepoints 0, 3, and 7 days post-injury (dpi), combined with Pax7^mTmG^ and age-matched cells from limb muscles reported in Tabula Muris Senis. Cells are colored by sample. B) Monocle pseudotime plot of aged MuSCs from the three timepoints post-injury displaying a branched trajectory. C) Alignment of the young and aged pseudotime trajectories via dynamic time warping, indicating differences in kinetics of the activation/differentiation programs of the two celltypes. An initial shift in aged cells at the start of the trajectory (i) indicated overexpression of senescence genes, followed by a large alignment region with rapid change in young MuSCs (ii). D) Scaled gene expression plots of young and along the pseudotime alignment trajectory. Igfbp5 showed strong differential expression between young and aged in region (i) of the trajectory, whereas Fibronectin and MyoG showed strong differential expression between the cell types in region (ii). E) Volcano plot displaying differential expression of genes from MyoG^hi^ MuSCs from state 3 after aged muscle injury (1,704 cells) to MyoG^hi^ MuSCs isolated from neurodegeneration (Sod1^-/-^) and age (155 cells). Significantly differentially expressed genes labeled in red highlight differences between the aged myogenic repair program adopted by young MuSCs after muscle injury and the activation program of Sod1^-/-^ and aged MuSCs.

Over the time points queried after muscle injury, we did not detect significant activation of markers associated with an NMJ-committed phenotype. To further assess the differences in expression between MuSCs that were upregulated in response to neuro-degeneration MyoG^+^/Dok7^+^ (which were primarily derived from uninjured Sod1^-/-^ and aged muscle) with young MuSCs that were MyoG^+^ after BaCl_2_ injury, differential expression analysis was performed on the two populations (Fig. 4f). MyoG^+^ MuSCs after muscle injury were strongly enriched for myogenic contractile proteins (Tnnt3, Tpm1, Eno3, Tnnc2, Lgals3), which was in contrast to markers from activation through neuro-degeneration (MyoG^+^/Dok7^+^) such as ubiquitin-proteasome (UPS) genes (Ubc, Uba52, Zfand5^52^) as well as neuro-regenerative transcripts (Nr4a1, Cux1, Btg1, Btg2). These results show a divergence in the gene expression program between the healthy myogenic repair adopted by MuSCs after muscle injury to the nervous rescue program presumed aimed at innervation rescue in from Sod1^-/-^ and aged MuSCs.

### Aged Muscle Stem Cells Displays Shifts in Regenerative Continuum Compared to Young After Muscle Injury but Do Not Acquire an NMJ-Committed Phenotype

Given the number of NMJ-committed MuSCs in aged muscle before injury was increased relative to young, we next determined if this population increased after muscle injury. Aged MuSCs were isolated after BaCl_2_ injury from the same time periods (3 and 7 dpi) as above using FACS and scRNA-Seq was performed, capturing 20,466 cells after filtering. UMAP clustering of aged MuSCs with P7mTmG datasets and cells isolated from age-matched limb muscles in Tabula Muris Senis once again yielded a strong MuSC cluster as well as small percentages of cells associated with neuronal-related, endothelial, fibroblast, and immune cell types (Fig 5a, Supp. Fig. 4a-b). Overlaying the expression of myogenic TFs onto UMAP plots showed recovery of different states in a similar temporal progression as young MuSCs (Fig. 5a, Supp. Fig. 4a). These results show that despite the MuSC heterogeneity before and after injury, separation from the NMJ-associated cluster was observed in a similar fashion to young MuSCs. Similar to young MuSCs, a decrease in the percentage of FSMs that overlapped with the NMJ-associated cluster (Supp. Fig. 4c) was observed for 3 and 7 dpi, respectively.

To further understand how single aged MuSCs varied in their regenerative trajectory compared to young, the cells were ordered in pseudotime (Fig. 5b). The trajectory revealed aged MuSCs displayed a similar course as young MuSCs but with differences in kinetics (Supp. Fig. 6d-e). To quantitatively understand the temporal differences, the young MuSC pseudotime trajectory was contrasted to the aged MuSC pseudotime trajectory using dynamic time warping^53^. Alignment of the two trajectories showed multiple off-diagonal alignments, with a large shift in aged cells at the start of the trajectory (Fig. 5c). Analysis of the genes driving the shift in aged MuSCs revealed increases in expression of genes associated with cellular senescence such as Igfbp5^54^. After the initial shift by aged MuSCs, the pseudotime trajectory was directed by young MuSCs via increases in expression of genes associated with matrix deposition such as Fibronectin (Fn1) followed by increases in MyoG (Fig. 5d). Summing these results shows young MuSCs initiate their regenerative differentiation and deposition of regenerative matrix with faster kinetics.

We next set to test if the differences in expression between aged MuSCs undergoing muscle repair (MyoG^+^) were distinct from MuSCs responding to neuro-degeneration (MyoG^+^/Dok-7^+^). Aged MuSCs undergoing muscle regeneration were observed to display increased expression of myogenic differentiation genes (Des, Vim, Lgals1, Tpm1) similar to young MuSCs (Fig. 5e). In contrast, MuSCs from neuro-degenerative models overexpressed transcripts associated with neuro-regeneration (Nr4a1, Cux1, Lmo4) and genes associated with stress response (Atf3, Junb, Fos, Egr1, Ier2). Many of the UPS genes that were enriched in the neurodegenerative population when compared to young MuSCs undergoing muscle regeneration were not differentially expressed comparing the two populations.

## Discussion

The development and enhancement in sensitivities of genomic technologies have advanced our understanding of *in vivo* stem cell actions. Profiling purified populations of MuSCs from different age groups through the regenerative process and neurodegenerative models enabled high-resolution molecular definition of stem cell states. Since the discovery of MuSCs, the population has been shown to possess subsets that proliferate at different rates and display variations in regenerative potential^3,55^. The heterogeneity in these dynamics can be ascribed to different gene expression programs^56^, the origins of which have remained elusive^57^.

The neuro-muscular synapse has long been recognized to be controlled through bi-directional communication between MNs and synaptic myonuclei^41,58^. We show here that this communication circuitry extends to a subset of MuSCs that enact a compensatory response when this cross-talk becomes broken. Previously, genetic ablation of MuSCs have been shown to impair NMJ regeneration associated with loss of sub-synaptic myonuclei^17,18^ after SNT, and aged NMJs display fewer sub-synaptic myonuclei coincident with age-related loss of MuSC number and function. Furthermore, MuSCs in denervated muscle fail to terminally differentiate^59^ and form smaller myotubes^60,61^ suggesting not only is the intricate interaction between MuSCs and MNs pivotal for regeneration but also to defend against nervous insults^62^ that result in muscle atrophy. Our analysis of gene expression of single MuSCs revealed changes in a subset of MuSCs through increases in expression of synaptic proteins such as Rapsyn and S100β. The increased synthesis of these molecules in subsets of MuSCs in our aged and Sod1^-/-^ models is consistent with previous observations that denervation induces expression of synaptic genes in muscle^39,40^. Our findings from SynTgSod1^-/-^ mice that displayed decreases in NMJ degeneration and reductions in the number of NMJ-committed MuSCs suggests MuSCs act in an anterograde manner in response to denervation. In further support of this model, we observed that MuSCs displayed focused engraftment into post-synaptic locations in the central muscle zone after denervation in young muscle. However, in aged muscle, MuSCs did not engraft but instead migrated towards NMJs^17^ and developed long filopodia. Summing these results together indicate MuSC-based replenishment of postsynaptic nuclei is compromised in aging and a subset of NMJ-committed progenitors remain, in agreement with our scRNA-Seq results. This is further supported by previous work demonstrating aging is associated with a loss of MuSC derived contributions in the vicinity of NMJs, a reduction in post-synaptic myonuclei, and accelerated NMJ degeneration upon MuSC depletion^17^. The attractive interaction between MuSCs and denervation requires further study, but development of filopodia has previously been observed for MuSCs^14,63^ and may facilitate transport of signaling ligands^64^ or local changes in cadherin signaling^65^ that is critical for synaptic patterning^66^.

The control of stem cell fate observed in aged and Sod1^-/-^ models towards an NMJ-committed phenotype may be influenced at several levels. First, aged myofibers undergoing atrophy from NMJ deterioration have been shown to accumulate reactive oxygen species^67^, which in turn signal to MuSCs to activate p38-MAPK signaling^68^. The gain of oxidative stress and associated loss of quiescence and activation is concordant with our observations. Next, while the role of Wnt signaling in synaptic formation is unclear^69^, our results suggest MuSCs may use this pathway to initiate an expression program prior to differentiation that activates AChR pre-patterning genes for subsequent synaptogenesis. In line with this, MuSCs have been shown to emit the Wnt7a ligand^70^ during differentiation, which also promotes axonal remodeling and synaptic differentiation^71^, and our results of Rapsyn expression increasing with upregulation of Wnt signaling agrees with these results^72^. Last, aging has been observed to induce dysregulation of calcium (Ca^2+^) homeostasis, and calcium-sensitive proteins such as S100β were observed to be upregulated in aged and Sod1^-/-^ MuSCs. The mechanism of dysregulated Ca^2+^ signaling in aged MuSCs remains unresolved, but Ca^2+^ influx through mechano-sensitive cation channels or voltage-gated Ca^2+^ channels may contribute^73^. Given MuSCs position on top of the sarcolemma, which becomes more permeable with age, myofiber membrane depolarization and associated changes in Ca^2+^ flux may contribute to NMJ gene expression^74^. Additionally, Ca^2+^-induced calmodulin-dependent protein kinase (CaMK) signaling has been observed to activates expression of genes associated with AChR clustering and NMJ formation such as Rapsyn^75^. Taken together, these data suggest NMJ deterioration, as observed in aging, engenders pathway activation in MuSCs that supports reinnervation and pre-patterning behavior.

The heterogeneity of MuSCs during regeneration has been shown to operate along a continuum^3, 76, 77^ and consistent with this view, we viewed waves of expression changes after injury in MuSCs. Organizing the MuSC expression profiles in pseudotime and overlaying myogenic transcription factor expression displayed Pax7^hi^ MuSCs were enriched at the root of the trajectory while MyoG^hi^ MuSCs were clustered towards the end of the trajectory. This pattern was conserved between young and aged MuSCs but the kinetics of these expression patterns and magnitude varied. In contrast to activation by neuro-degeneration, the activation trajectory of both young and aged MuSCs from muscle injury was distinct and induced transcription of different gene sets. These results show gene transcription in MuSCs in response to different injuries confer different regenerative trajectories.

The comprehensive approaches used in the present study begin to provide insights into the mechanisms and functional consequences of MuSC heterogeneity with age and provide a resource for further understanding of non-regenerative muscle diseases, anti-aging models and severe forms of trauma such as volumetric muscle loss^78^.

## Acknowledgments

The authors thank Daniel Vincenz for assistance with artwork, and the University of Michigan DNA Sequencing Core for assistance with single cell sequencing library preparation. The authors also thank Alex Shalek for critical advice regarding the single cell sequencing and data processing, Josh Welch for insights into bioinformatics analysis, and members of the Aguilar and Brooks laboratories. Research reported in this publication was partially supported by the National Institute of Arthritis and Musculoskeletal and Skin Diseases of the National Institutes of Health under Award Number P30 AR069620 (CAA, SVB), the Breast Cancer Research Foundation (PJU, SDM), the 3M Foundation (CAA), American Federation for Aging Research Grant for Junior Faculty (CAA), the University of Michigan Geriatrics Center and National Institute of Aging under award number P30 AG024824 (CAA, SVB), the University of Michigan Biomedical Engineering Department (CAA), the Department of Defense and Congressionally Directed Medical Research Program W81XWH18SBAA1-12579992 (CAA, YCJ), the National Institute on Aging P01 AG051442 (SVB), and National Institute on Aging R01 AG051456 (JVC). The content is solely the responsibility of the authors and does not necessarily represent the official views of the National Institutes of Health.

## Accession Code

GEO: 121589

## Author contributions

J.L., M.M., J.C.M., W.L., L.A.B., J.F.M., J.J.C., K.D.S., B.D.L., J.V.C., Y.C.J., and C.A.A. performed experiments. P.J.U., J.L., S.J.K., and C.A.A. analyzed data. S.V.B and C.A.A. designed the experiments and C.A.A. wrote the manuscript with additions from other authors.

## KEY RESOURCES TABLE

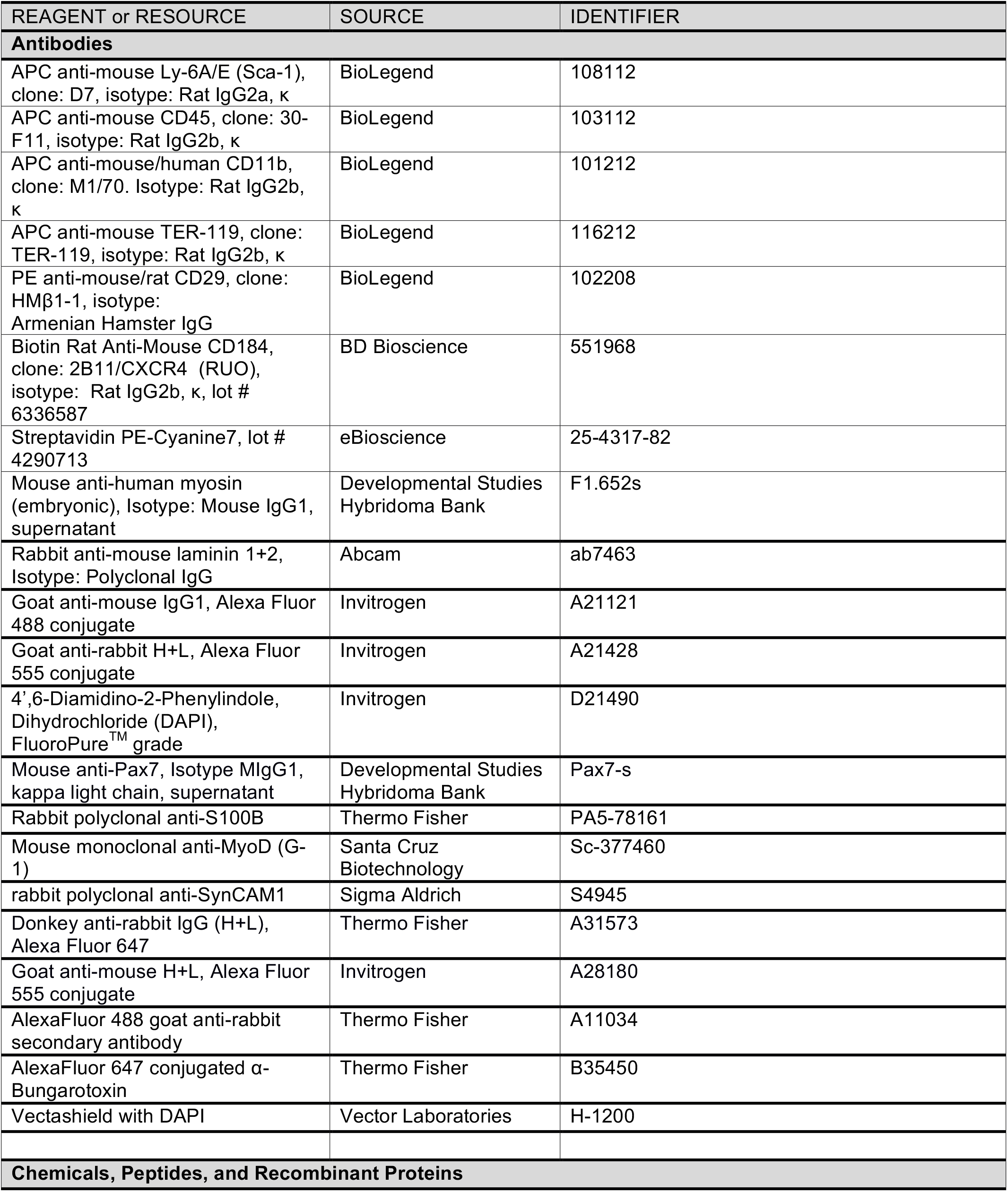
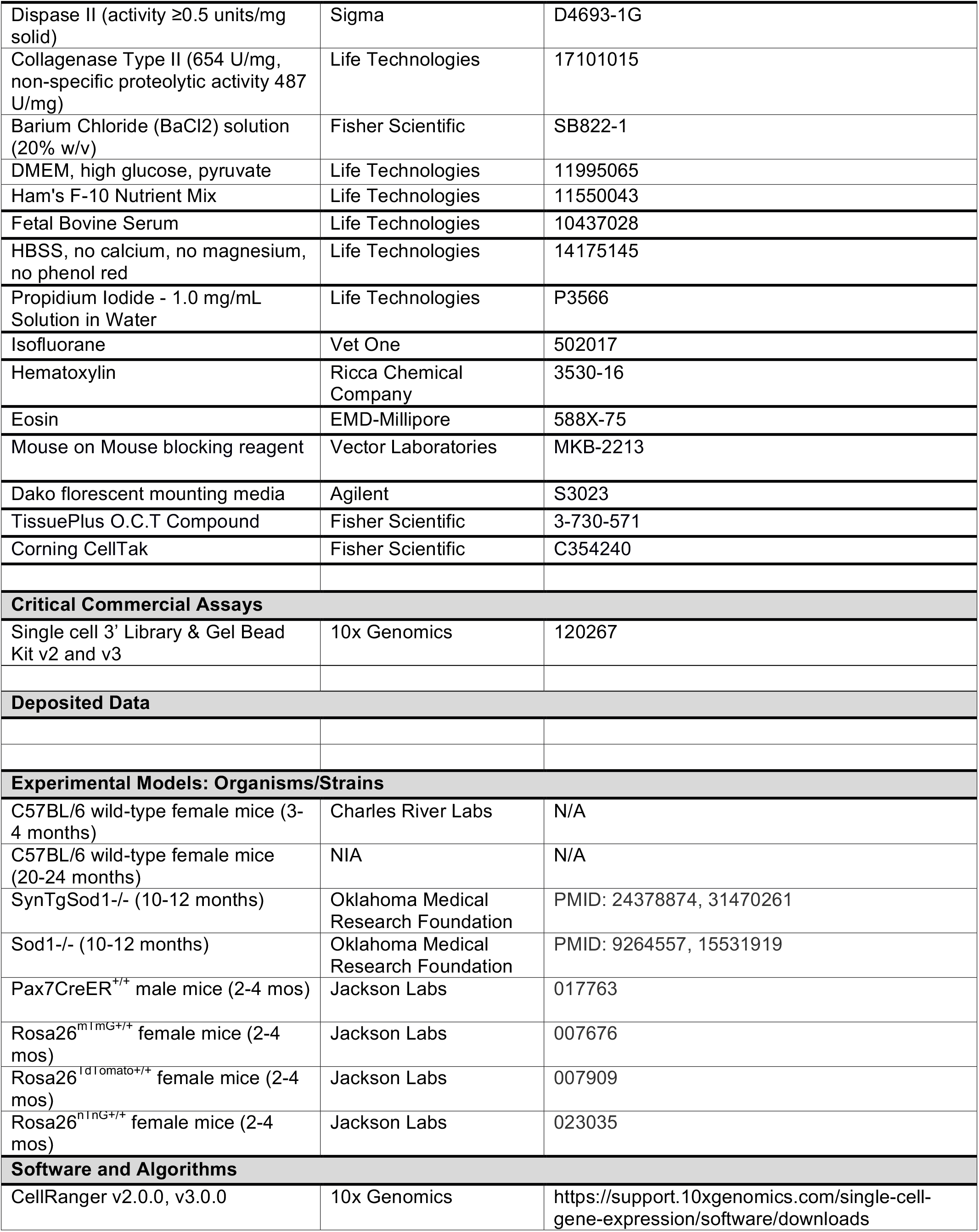
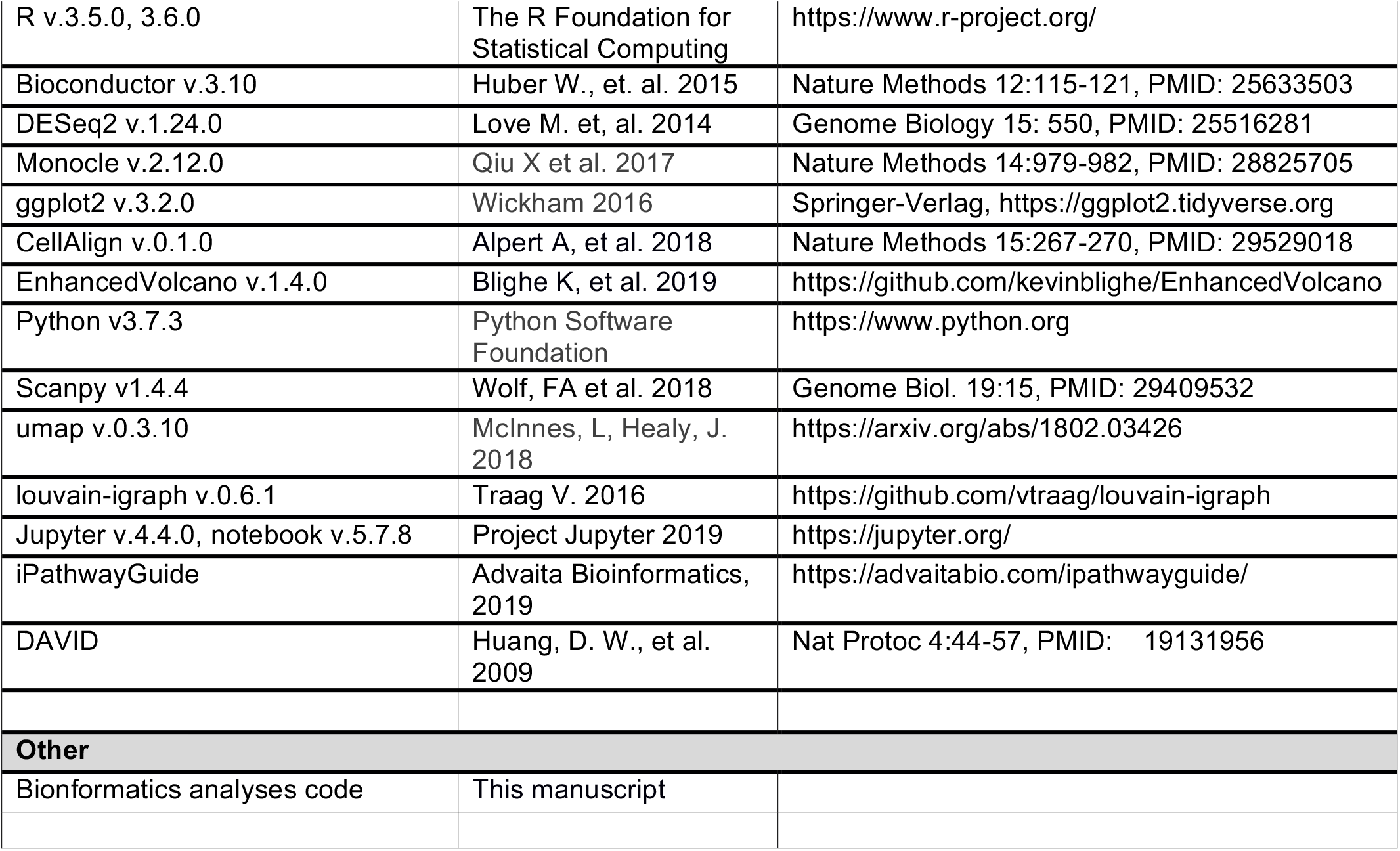

## Materials and Methods

### Animal and Injury Model

C57BL/6 wild-type female mice were obtained from Charles River Breeding Laboratories, the National Institute on Aging, or from a breeding colony at the University of Michigan (UM). 2- to 3-month-old Pax7^CreER/+^ male mice were crossed with 2- to 3-month-old Rosa26^mTmG/+^ female mice to generate Pax7^CreER/+^-Rosa26^mTmG/+^ and genotyped. 2- to 3-month-old Pax7^CreER/+^ male mice were bred with mice carrying a loxP-flanked STOP cassette followed by *TdTomato* in the ROSA26 locus and genotyped. To activate GFP expression in the Pax7^CreER/+^-Rosa26^mTmG/+,^ Pax7^CreER/+^-Rosa26^nTnG/+^ mice, mice were administered tamoxifen through intraperitoneal injection five times (1x per day), and allowed to recover for 2-5 days. Pax7^CreER/+^-Rosa26^TdTomato/+^ mice were fed with tamoxifen chow for at least 2 weeks prior to euthanasia to induce TdTomato expression in Pax7^+^ cells through the Cre recombinase activity and recombination efficiency was measured at >90%. All mice were fed normal chow ad libitum and housed on a 12:12 hour light-dark cycle under UM veterinary staff supervision. All procedures were approved by the University Committee on the Use and Care of Animals at UM and were in accordance with the U.S. National Institute of Health (NIH). Young female mice (3-4 months) and aged female mice (20-24 months) were randomly assigned to one of five groups: uninjured, day 3, and day 7 injured (n=4 per group). To induce skeletal muscle injury, mice were first anesthetized with 2% isoflurane and administered a 1.2% barium chloride (BaCl_2_) solution injected intramuscularly into several points of the tibialis anterior and gastrocnemius muscles for a total of 80 μL per hindlimb.

### Satellite Cell Isolation via Fluorescence-Activated Cell Sorting

For tissue collection, mice were anesthetized with 3% isoflurane, then euthanized by cervical dislocation, bilateral pneumothorax and removal of the heart. Hind limb muscles (tibialis anterior and gastrocnemius) of control and experimental mice were quickly harvested using sterile surgical tools and placed in separate plastic petri dishes containing cold PBS. Using surgical scissors, muscle tissues were minced and transferred into 50 mL conical tubes containing 20 mL of digest solution (2.5 U/mL Dispase II and 0.2% [~5,500 U/mL] Collagenase Type II in DMEM media per mouse). Samples were incubated on a rocker placed in a 37°C incubator for 60 min with manual pipetting the solution up and down to break up tissue every 30 minutes using an FBS coated 10 mL serological pipette. Once the digestion was completed, 20 mL of F10 media containing 20% heat inactivated FBS was added into each sample to inactivate enzyme activity. The solution was then filtered through a 70 μm cell strainer into a new 50 mL conical tube and centrifuged again at 350xg for 5 min. The pellets were re-suspended in 6 mL of staining media (2% heat inactivated FBS in Hank’s Buffered Salt Solution - HBSS) and divided into separate FACS tubes. The FACS tubes were centrifuged at 350xg for 5 min and supernatants discarded. The cell pellets were then re-suspended in 200 μL of staining media and antibody cocktail containing Sca-1:APC (1:400), CD45:APC (1:400), CD11b:APC (1:400), Ter119:APC (1:400), CD29/β1-integrin:PE (1:200), and CD184/CXCR-4: BIOTIN (1:100) and incubated for 30 minutes on ice in the dark. Cells and antibodies were diluted in 3mL of staining solution, centrifuged at 350xg for 5 min, and supernatants discarded. Pellets were resuspended in 200uL staining solution containing PECy7:STREPTAVIDIN (1:100) and incubated on ice for 20 minutes in the dark. Again, samples were diluted in 3mL staining solution, centrifuged, supernatants discarded, and pellets re-suspended in 200uL staining buffer. Live cells were sorted from the suspension via addition of 1 μg of propidium iodide (PI) stain into each experimental sample and all samples were filtered through 70 μm cell strainers before the FACS. Cell sorting was done using a BD FACSAria III Cell Sorter (BD Biosciences, San Jose, CA) and APC negative, PE/PECy7 double-positive MuSCs were sorted into staining solution for immediate processing.

### Single-Cell mRNA Sequencing

Freshly isolated MuSCs were sorted into staining solution, enumerated by hemocytometer, and re-suspended into PBS. Cells were loaded into the 10x Genomics chromium single cell controller for each time point and age group were captured into nanoliter-scale gel bead-in-emulsions (GEMs). cDNAs were prepared using the single cell 3’ Protocol as per manufacturer’s instructions and sequenced on a NextSeq 500 instrument (Illumina) or NovaSeq instrument (Illumina) with 26 bases for read1 and 98 bases for read2.

### Histological and Immunostaining of Hindlimb Muscle

A separate cohort of mice was used for histological evaluation of skeletal muscle injury and regeneration. Young (3-4 months) and old (20-24 months) female mice (n=4 per group) received bilateral intramuscular injection of the tibialis anterior muscles with a 1.2% BaCL_2_ solution (40 μL per muscle) while under isoflurane anesthesia. At day 3 and 7 post BaCl_2_ injury, mice were euthanized via bilateral pneumothorax and cervical dislocation, and tibialis anterior muscles were rapidly dissected. Tibialis anterior muscles were embedded in optimum cutting temperature (OCT) compound (Tissue Plus, Fisher Scientific), rapidly frozen in isopentane, cooled on liquid nitrogen and stored at −80°C until further analysis.

Muscle tissue cross sections (10 μm) were cut in a cryostat at −20°, adhered to Superfrost Plus microscope slides (Fisher Scientific) and air dried at room temperature. Slides were fixed in 100% acetone for 10 minutes at −20°C, air dried at room temperature and stained with hematoxylin and eosin (H&E) using routine procedures. Sections for immunofluorescence staining were rehydrated in PBS for 5 minutes and blocked overnight at 4°C in M.O.M. blocking reagent. Sections were then incubated overnight at 4°C with a cocktail of primary antibodies against embryonic myosin (1:20 dilution) and laminin (1:200 dilution). The following morning, slides were washed three times for 5 minutes in PBS and incubated for 1 hour at room temperature with Goat Anti-Mouse IgG1 Alexa Fluor 488 (1:500) and Goat Anti-Rabbit Alexa Fluor 555 (1:500) secondary antibodies. Nuclei were counterstained with 4’,6-Diamidino-2-Phenylindole Dihydrochloride (DAPI) (1 μg/mL). Slides were washed three times for 5 minutes each in PBS and then mounted with coverslips using Dako Fluorescence mounting media. Fluorescent and bright field images were acquired using a Nikon A1 confocal and Olympus BX51 wide field microscope. Regenerating myofibers (central nuclei and/or eMyHC+ cytoplasm) were quantified in a single 20x field of view from the core of the injured region using Image J software. Differences between young and aged mice for regenerating myofibers and regenerating myofiber cross-sectional areas were determined by FIJI and student’s t-tests where statistical significance was set at *p* < 0.05.

### Immunostaining of Neuromuscular Junctions

To identify Pax7 expressing satellite cells, *Pax7^CreER/+^; R26LSL^TdTomato/+^* mice were used as described previously^1^. Extensor digitorum longus (EDL) and soleus muscles were excised and fixed for 2 hours at room temperature (4% paraformaldehyde in PBS). Unless otherwise indicated, all incubations were performed at room temperature. EDL muscles were teased into bundles of muscle fibers. Muscles were washed twice for 15 min. in PBS, incubated 15 min. with 100 mM glycine in PBS, and rinsed in PBS. After removal of the overlying connective tissue, muscles were permeabilized and blocked for 1 hr. using M.O.M. blocking reagent then in PBS containing 2% bovine serum albumin, 4% normal goat serum, and 0.5% Triton X-100 for another hour. To detect post-synaptic acetylcholine receptors (AChRs), muscles were stained with AlexaFluor 647 conjugated α-Bungarotoxin (1:200), first for 1 hour at room temperature and then overnight at 4°C. After three 30 min. washes in PBS, muscles were rinsed in PBS, fixed 30 min. with 1% formaldehyde in PBS, washed three times for 10 min. in PBS, and flat-mounted in Vectashield with DAPI (Vector Laboratories, Cat# H-1200). Images were acquired using a Zeiss Axio Observer Z1 or LSM 780 confocal microscope.

### Immunostaining of S100B, Pax7 and MyoD

Clear 8-well coverslides were coated with 22.4ug/mL CellTak in PBS for 20 minutes at room temperature, followed by three quick rinses with distilled water. FACS-enriched MuSCs were then seeded at a density of 10,000 cells/well and allowed to adhere for 45 minutes in staining buffer at room temperature. After adhering to the cover slide, staining buffer was aspirated and cells were fixed with 4% paraformaldehyde in PBS at room temperature for 20 minutes. Paraformaldehyde was aspirated and after three quick washes with PBS, cells were permeabilized with 0.1% TritonX-100 and blocked with 1% BSA, 0.1% Tween-20 and 22.52mg/mL glycine in PBS. After blocking, cells were incubated with primary antibodies (1:500 dilution of anti-S100B, 1:10 dilution of anti-Pax7, 1:200 dilution of anti-MyoD) overnight at 4°C followed by secondary antibodies (1:300 dilution of AF555 anti-mouse, 1:300 dilution of AF647 anti-rabbit) overnight at 4°C. Nuclei were stained with Hoechst 33342 (1.5ug/mL) in PBS for 1 minute at RT. Immunolabeled cells were imaged on a Zeiss epifluorescent microscope using a 10X objective. Single and double-positive cells were manually counted. Statistical comparison between groups was performed using a twosided, two sample student’s t-test assuming equal variances. P-values below 0.05 were considered significant.

### Sciatic Nerve Transection (SNT)

Mice were anesthetized with intraperitoneal injections of ketamine (110 mg/kg) and xylazine (10 mg/kg). The hindquarter was then carefully shaved and depilation completed with generic Nair hair removal cream prior to skin cleansing with gauze. The skin was incised 1 mm posterior and parallel to the femur, and the biceps femoris was bluntly split to expose the sciatic nerve. 1–2 mm sciatic nerve was then transected 5 mm proximal to its trifurcation, followed with realignment of the distal and proximal nerve ends and closure of the muscle with wound clips (Autoclip, BD Clay Adams, Franklin Lakes, NJ). Mice were given analgesic (0.5–1.0 mg/kg buprenorphine) and allowed to recover on a heating pad. Sham surgery was performed on the contralateral leg where procedures were performed without nerve transection.

### Single Myofiber Analysis

For nGFP localization, myofibers were purified by conventional collagenase digestion and trituration with fire polished glass pipets as previously described^2^. Briefly, the EDL muscle was dissected, rinsed in Dulbecco’s phosphate-buffered saline (PBS), put into a 1.5 ml Eppendorf tube containing 1 ml 0.1% type I collagenase (Invitrogen) and 0.1% type II collagenase (Invitrogen) in Dulbecco’s modified Eagles medium (DMEM, Sigma–Aldrich, St. Louis, MO), incubated in a shaker water bath at 37°C for 75 min and gently mixed by inversion periodically. Following digestion, the muscle was transferred to 100 mm Å~ 15 mm plastic petri dishes containing 10 ml of plating media (10% horse serum in DMEM) using fire-polished-tip Pasteur pipettes. Under a stereo dissecting microscope, single myofibers were released by gently triturating the EDL with a series of modified Pasteur pipettes that varied in tip diameter to accommodate the progressive decrease in muscle trunk size. Inseparable fibers and debris were removed. Purified single myofibers were fixed with 4% PFA for 3 min, washed with PBS and transferred to 5 ml polystyrene cell collection tubes for nGFP, Pax7 and Btx IF.

### scRNA-Seq Data Processing and Analysis

CellRanger v2.0 or v3.1 (10x Genomics) was used to process raw data. The CellRanger workflow aligns sequencing reads to the mm10 transcriptome using the STAR aligner^3^ and exports count data. The CellRanger count command was run with default parameters with the exception of the -- expect-cells parameter which was set at 10000. Filtered feature barcode data from all samples in HDF5 matrix format, both locally generated datasets as well as public Tabula Muris (TM)^4^ and Tabula Muris Senis (TMS) datasets^5^ were imported directly into Python3 ScanPy^6^ Anndata objects using the read_10x_mtx import method. Two base quality filters were applied, retaining only cells with a minimum of 300 genes expressed and genes expressed in at least five cells. All cell counts provided in the main manuscript text are post-filtering counts. Fifteen datasets were generated in this manner: C57BL/6 wild-type female mice both young (3-4 mo.) and aged (20-24 mo.) datasets at timepoint 0, and days 3 and 7 post-injury (6 datasets); P7^mTmG^ datasets (2); Sod1^-/-^ and SynTgSod1^-/-^ rescue mouse datasets (2); public TM (1) and TMS (4) datasets. Datasets were aggregated into the four collections discussed in this manuscript using the ScanPy/anndata concatenate method: the ‘uninjured’ collection (Fig. 1) composed of young, aged timepoint 0, the Pax7 data sets, and the TM and TMS datasets; the ‘neurodegenerative’ collection (Fig. 3) using young, aged timepoint 0, the Sod1 KO and rescue sets, and the TM and TMS datasets; the ‘young’ collection (Fig. 5) composed of young mice at timepoints 0, 3, and 7 post-injury, the Pax7+ cells, and the TM dataset; and the ‘aged’ collection (Fig. 6) using aged mice at timepoints 0, 3, and 7 post-injury, the P7mTmG cells, and the TMS dataset. We evaluated batch correction methods but did not utilize batch correction, as each dataset represented a distinct biological condition, with the exception of the Pax7^mTmG^ datasets (two separate samples) and the public Tabula Muris Senis data (four datasets). These data clustered quite closely as profiled in dimension reduction maps (Supp. Fig. 1E, for example). A small batch effect can be discerned between these replicate datasets, but they effect appeared intra-cluster rather than inter-cluster.

Mitochondrial gene fraction was calculated for cells in the aggregated datasets and cells with >15% mitochondrial genes were removed. Ribosomal genes were also filtered. Cells with abnormally large UMI counts were filtered out based on QC plotting to remove possible doublets; the threshold varied by dataset but ranged from 20,000 to 70,000 counts per cell. Aggregated datasets were then per-cell normalized using the ScanPy pp.normalize_total method, which size-corrects each cell by total counts over all genes. A logX + 1 transformation of count data was generated for downstream analysis, retaining a version of the raw counts data for subsequent calculations as well, noted below. A dispersion estimate was performed on genes using the ScanPy pp.highly_variable_genes method on the log-transformed data. Prior to dimension reduction, variation due to mitochondrial fraction and total counts was removed via linear regression using the pp.regress_out method.

Dimension reduction was performed on each dataset using principal component analysis as implemented in the ScanPy tl.pca method using the ARPACK implementation via SciPy/Scikit-learn and otherwise default parameters. The number of components to utilize for subsequent neighborhood graph calculation was determined by generating an elbow plot, resulting in either 40 or 45 components. The neighborhood graph was then computed using the pp.neighbors method^7^. 2D embeddings were then calculated using the tl.umap function with default parameters, and clustering into subgroups performed using the tl.louvain method with various resolutions, with resolution 0.5 most commonly utilized as optimal to capture the underlying biology based on inspection of resulting marker gene lists for resulting clusters. Marker gene selection was performed using the ScanPy tl.rank_genes_groups method, utilizing a Wilcoxon rank sum statistic and otherwise default parameters. For several additional analysis discussed in the manuscript, subsets of cells were selected from each collection using manual python matrix operations on the ScanPy Anndata object, choosing cells with specific cluster identifications or properties such as the expression of particular genes (e.g. ‘Myog’), in some cases followed by re-clustering of the subsets using the tl.louvain method. Differential gene expression measurements of the Sod1-/- mice vs young and aged samples were calculated using DESeq2 using a direct ‘condition a vs b’ model using only genes expressed in both datasets. Volcano plots were generated using the R package EnhancedVolcano^8^.

### Quiescence score calculation

A quiescence score calculation was implemented by calculating the total fraction of the counts in each cell derived from quiescence-associated genes. A list of 45 quiescent genes was utilized, 40 of which were expressed in our datasets after filtering (Supp. Table 1). The raw counts associated with each of these genes were summed, and the sum divided by the total UMI count across all genes in each cell to provide a normalized sum score.

### Monocle Cell pseudotime trajectory estimation

Cell pseudotime differentiation trajectories were calculated on the MuSC cell cluster subsets of the ‘young’ and ‘aged’ cell collections using the Monocle R package^9^ (version 2). Raw count data were exported from Python/Scanpy AnnData objects as matrices, as well as cell and gene specific information from AnnData.obs and AnnData.var objects, respectively, as R AnnotatedDataFrame objects. These objects were used to create monocle CellDataSet (CDS) objects. Size factors and dispersions were calculated on the CDS objects using methods available in the DESeq2 R package^10^. ‘Ordering’ genes differentially expressed as a function of time point were detected using the Monocle differentialGeneTest method using the model formula ~day, where ‘day’ is a value assigned to each cell based on time post-injury (‘0’, ‘3’, or ‘7’); ordering genes were selected based on a q-value < 0.005. Cells were mapped to a 2D plane using the DDRTree method^11^, and ordered in pseudotime using the orderCells function with default parameters. Genes that differ between the three branches were identified via the Monocle BEAM method^12^ utilized to generate the split heatmaps.

### CellAlign trajectory alignment

Normalized gene expression cell matrices and the inferred pseudotime values for each cell were extracted from the Monocle2 CellDataSet objects for young and aged samples. The data were interpolated to construct a 200 point equally spaced pseudotime trajectory space using the CellAlign interWeights method and scaled using the scaleInterpolate method^13^. Finally, the interpolated/scaled trajectories for Young and Aged datasets were aligned using CellAlign’s globalAlign method using the default Euclidean distance metric and the ‘symetric2’ step pattern. Plotting of the alignment was perfomed using the plotAlign method.

---

**Supplemental Table 1**. List of quiescent genes used to calculate quiescent score.

**quiescence**

Pax3

Pax7

Foxo3

Ezh1

Itm2a

Zbtb20

Notch1

Notch2

Notch3

Hoxa9

Hes1

Heyl

Pdk4

Trp63

Rara

Rarb

Rarg

Calcr

Col5a1

Sox7

Sox18

Meox1

Meox2

Sdc4

Cd34

Rbpj

Cav1

Cxcr4

Itgb1

Itga7

Tenm4

Vcam1

Emd

Spry1

Fzd7

Cebpb

Pde4b

Zfp36

Gpx3

Fgl2

## References

1. Yin, H. et al. MicroRNA-133 controls brown adipose determination in skeletal muscle satellite cells by targeting Prdm16. Cell Metab 17, 210–224 (2013).

2. Zammit, P.S. et al. Muscle satellite cells adopt divergent fates : a mechanism for self-renewal? The Journal of Cell Biology 166, 347–357 (2004).

3. Dobin, A. et al. STAR: ultrafast universal RNA-seq aligner. Bioinformatics (Oxford, England) 29, 15–21 (2013).

4. Schaum, N. et al. Single-cell transcriptomics of 20 mouse organs creates a Tabula Muris. Nature 562, 367–372 (2018).

5. Pisco, A.O. et al. A Single Cell Transcriptomic Atlas Characterizes Aging Tissues in the Mouse. bioRxiv, 661728 (2019).

6. Wolf, F.A., Angerer, P. & Theis, F.J. SCANPY: large-scale single-cell gene expression data analysis. Genome Biol 19, 15 (2018).

7. McInnes & Healy UMAP: Uniform manifold approximation and projection for dimension reduction. arXiv 1802.03426 (2018).

8. Blighe, K., Rana, S. & Lewis, M. (https://github.com/kevinblighe.; 2018).

9. Qiu, X. et al. Reversed graph embedding resolves complex single-cell trajectories. Nat Methods 14, 979–982 (2017).

10. Love, M.I., Huber, W. & Anders, S. Moderated estimation of fold change and dispersion for RNA-seq data with DESeq2. Genome biology 15, 550–550 (2014).

11. Mao, Q., Wang, L., Goodison, S. & Sun, Y. in Proceedings of the 21th ACM SIGKDD International Conference on Knowledge Discovery and Data Mining 765–774 (ACM, Sydney, NSW, Australia; 2015).

12. Qiu, X. et al. Single-cell mRNA quantification and differential analysis with Census. Nat Methods 14, 309–315 (2017).

13. Alpert, A., Moore, L.S., Dubovik, T. & Shen-Orr, S.S. Alignment of single-cell trajectories to compare cellular expression dynamics. Nat Methods 15, 267–270 (2018).

## References

1. Marcell, T. Sarcopenia: causes, consequences and preventions. J. Gerontol. A. 58, M911-M916 (2003).

2. Wang, Y.X., Rudnicki, M.A. Satellite cells, the engines of muscle repair. Nature Rev. Molec. Cell Biol. 13, 127–133 (2012).

3. Porpiglia, E. et al. High-resolution myogenic lineage mapping by single-cell mass cytometry. Nature Cell Biol. 19, 558–567 (2017).

4. Blau, H.M., Cosgrove, B.D., Ho, A.T.V. The central role of muscle stem cells in regenerative failure with aging. Nature Med. 21, 854–862 (2015).

5. Yin, H.F., Price, F., Rudnicki, M.A. Satellite cells and the muscle stem cell niche. Physiological Reviews 93, 23–67 (2013).

6. Almada, A.E, Wagers, A.J. Molecular circuitry of stem cell fate in skeletal muscle regeneration, ageing and disease. Nature Rev. Molec. Cell Biol. 17, 267–279 (2016).

7. Goodell, M.A, Rando, T.A. Stem cells and healthy aging. Science 350, 1199–1204 (2015).

8. Schultz, E., Lipton, B.H. Skeletal muscle satellite cells: changes in proliferation potential as a function of age. Mech. Aging Dev. 20, 377–383 (1982).

9. Collins, C.A., Olsen, I., Zammit, P.S., Heslop, L., Petrie, A., Partridge, T.A., Morgan, J.E. Stem cell function, self-renewal, and behavioral heterogeneity of cells from the adult muscle satellite cell niche. Cell 122, 289–301 (2005).

10. Daya, J. et al. Biophysical and biomolecular determination of cellular age in humans. Nature Biomed. Engin. 1, 0093 (2017).

11. Tierney, M.T., Stec, M.J., Rulands, S., Simons, B.D., Sacco, A. Muscle stem cells exhibit distinct clonal dynamics in response to tissue repair and homeostatic aging. Cell Stem Cell 22, 119–127 (2018).

12. Gawad, C., Koh, W., Quake, S.R. Single-cell genome sequencing: current state of the science. Nature Rev. Genet. 17, 175–188 (2016).

13. Scharner, J., Zammit, P.S. The muscle satellite cell at 50: the formative years. Skeletal Muscle 1, 28 (2011).

14. Verma, M. et al. Muscle satellite cell cross-talk with a vascular niche maintains quiescence via VEGF and Notch signaling. Cell Stem Cell 23, 530–543 (2018).

15. Christov, C. et al. Muscle satellite cells and endothelial cells: close neighbors and privileged partners. Mol. Biol. Cell 18, 1397–1409 (2007).

16. Kostallari, E., Baba-Amer, Y., Alonso-Martin, S., Ngoh, P., Relaix, F., Lafuste, P., Gherardi, R.K. Pericytes in the myovascular niche promote post-natal myofiber growth and satellite cell quiescence. Develop. 142, 1242–1253 (2015).

17. Liu, W., et al. Loss of adult skeletal muscle stem cells drives age-related neuromuscular junction degeneration. Elife 6, e26464 (2017).

18. Liu, W., Wei-LaPierre, L., Klose, A., Dirksen, R.T., Chakkalakal, J.V. Inducible depletion of adult skeletal muscle stem cells impairs the regeneration of neuromuscular junctions. eLife 4, e09221 (2015).

19. Murphy, M.M., Lawson, J.A., Mathew, S.J., Hutcheson, D.A., Kardon, G. Satellite cells, connective tissue fibroblasts and their interactions are crucial for muscle regeneration. Develop. 138, 3625–3637 (2011).

20. Jang, Y.C., Van Remmen, H. Age-associated alterations of the neuromuscular junction. Exp. Geront. 46, 193–198 (2011).

21. Cerletti, M., Jurga, S., Witczak, C.A., Hirshman, M.F., Shadrach, J.L., Goodyear, L.J., Wagers, A.J. Highly efficient, functional engraftment of skeletal muscle stem cells in dystrophic muscles. Cell 134, 37–47 (2008).

22. Aguilar, C.A. et al. Transcriptional and chromatin dynamics of muscle regeneration after severe trauma. Stem Cell Rep. 7, 983–997 (2016).

23. Keefe, A.C., Lawson, J.A., Flygare, S.D., Fox, Z.D., Colasanto, M.P., Mathew, S.J., Yandell, M., Kardon, G. Muscle stem cells contribute to myofibers in sedentary adult mice. Nature Commun. 6, 7087 (2015).

24. Zheng, G.X.Y. et al. Massively parallel digital transcriptional profiling of single cells. Nature Commun. 8, 141049 (2017).

25. Becht, E., McInnes, L., Healy, J., Dutertre, C.A., Kwok, I.W.h., Ng, L.G., Ginhoux, F., Newell, E.W. Dimensionality reduction for visualizing single-cell data using UMAP. Nature Biotech. 37, 38–44 (2019).

26. The Tabula Muris Consortium. Single-cell transcriptomics of 20 mouse organs creates a Tabula Muris. Nature 562, 367–372 (2018).

27. Fukada, S.I., et al. Molecular signature of quiescent stem cells in adult skeletal muscle. Stem Cells 25, 2448–2459 (2007).

28. Lala-Tabbert, N., AlSudais, H., Marchildon, F., Fu, D., Wiper-Bergeron, N. CCAAT/enhancer binding protein beta is required for satellite cell self-renewal. Skeletal Muscle 6:40 (2016).

29. Sorci, G., Riuzzi, F., Arcuri, C., Tubaro, C., Bianchi, R., Giambanco, I., Donato, R. S100B in tissue development, repair and regeneration. World J. Biol. Chem. 4, 1–12 (2013).

30. Tanabe, Y., Fujita, E., Hayashi, Y.K., Zhu, X., Lubbert, H., Mezaki, Y., Senoo, H., Momoi, T. Synaptic adhesion molecules in Cadm family at the neuromuscular junction. Cell Biol. Int’l. 37, 731–736 (2013).

31. Biederer, T., Sara, Y., Mozhayeva, M., Atasoy, D., Liu, X., Kavalali, E.T., Sudhof, T. SynCAM, a synaptic adhesion molecules that drives synapse assembly synapse assembly and disassembly. Science 297, 1525–1531 (2002).

32. Goda, Y., Davis, G.W. Mechanisms of synapse assembly and disassembly. Neuron 40, 243–264 (2003).

33. Zhong, Z., Ohnmacht, J., Reimer, M.M., Bach, I., Becker, T., Becker, C.G. Chondrolectin mediates growth cone interactions of motor axons with an intermediate target. J. Neurosci. 32, 4426–4439 (2012).

34. Zhang, Z., Lotti, F., Dittmar, K., Younis, I., Wan, L., Kasim, M., Dreyfuss, G. SMN deficiency causes tissue-specific perturbations in the repertoire of snRNAs and widespread defects in splicing. Cell 133, 585–600 (2008).

35. Aare, S., Spendiff, S., Vuda, M., Elkrief, D., Perez, A., Wu, Q., Mayaki, D., Hussain, S.N.A., Hettwer, S., Hepple, R.T. Failed reinnervation in aging skeletal muscle. Skeletal Muscle 6, 29 (2016).

36. Zammit, P.S., Relaix, F., Nagata, Y., Ruiz, A.P., Collins, C.A., Partridge, T.A., Beauchamp, J.R. Pax7 and myogenic progression in skeletal muscle satellite cells. J. Cell Sci. 119, 1824–1832 (2006).

37. Jang, Y.C. et al. Increased superoxide in vivo accelerates age-associated muscle atrophy through mitochondrial dysfunction and neuromuscular junction degeneration. FASEB J. 24 1376–1390 (2010).

38. Parisi, A., Lacour, F., Giordani, L., Colnot, S., Maire, P., Le Grand, F. APC is required for muscle stem cell proliferation and skeletal muscle tissue repair. J. Cell Biol. 210, 717–726 (2015).

39. Macpherson, P.C.D., Wang, X., Goldman, D. Myogenin regulates denervation-dependent muscle atrophy in mouse soleus muscle. J. Cell. Biochem. 112, 2149–2159 (2011).

40. Moresi, V., et al. Myogenin and class II HDACs control neurogenic muscle atrophy by inducing E3 ubiquitin ligases. Cell 143, 35–45 (2010).

41. Li, L., Xiong, W.C., Mei, L. Neuromuscular junction formation, aging and disorders. Annu. Rev. Physiol. 80, 159–188 (2018).

42. Sakellariou, G.K. et al. Neuron-specific expression of CuZnSOD prevents the loss of muscle mass and function that occurs in homozygous CuZnSOD-knockout mice. FASEB J. 28 1666–1681 (2014).

43. Alonso-Martin, S. et al. Gene expression profiling of muscle stem cells identifies novel regulators of postnatal myogenesis. Front. Cell Develop. Biol. 4:57 (2016).

44. Pallafacchina, G. et al. An adult tissue-specific stem cell in its niche: a gene profiling analysis of in vivo quiescent and activated muscle satellite cells. Stem Cell Res. 4, 77–91 (2010).

45. Kelly, A.M. Perisynaptic satellite cells in the developing and mature rat soleus muscle. Anat. Rec. 190, 891–903 (1978).

46. Dedkov, E.I., Borisov, A.B., Wernig, A., Carlson, B.M. Aging of skeletal muscle does not affect the response of satellite cells to denervation. J. Histochem. & Cytochem. 51, 853–863 (2003).

47. Trapnell, C., Cacchiarelli, D., Grimsby, J., Pokharel, P., Li, S., Morse, M., Lennon, N.J., Livak, K.J., Mikkelsen, T.S., Rinn, J.L. The dynamics and regulators of cell fate decisions are revealed by pseudotemporal ordering of single cells. Nature Biotech. 32, 381–386 (2014).

48. Kops, G.J.P.L. et al. Forkhead transcription factor FOXO3a protects quiescent cells from oxidative stress. Nature 419, 316–321 (2002).

49. Agudo, J., Park, E.S., Rose, S.A., Alibo, E., Sweeney, R., Dhainaut, M., Kobayashi, K.S., Sachidanandam, R., Baccarini, A., Merad, M., Brown, B.D. Quiescent tissue stem cells evade immune surveillance. Immunity 48, 271–285 (2018).

50. Baghdadi, M.B., Castel, D., Machado, L., Fukada, S., Birk, D.E., Relaix, F., Tajbaksh, S., Mourikis, P. Reciprocal signaling by Notch-collagen V-CALCR retains muscle stem cells in their niche. Nature 514, 714–718 (2018).

51. Zhang, T., Gunther, S., Looso, M., Kunne, C., Kruger, M., Kim J., Zhou, Y., Braun, T. Prmt5 is a regulator of muscle stem cell expansion in adult mice. Nature Comm. 6, 7140 (2014).

52. Lee, D., Takayam, S., Goldberg, A.L. Zfand5/Znf216 is an activator of the 26S proteasome that stimulates overall protein degradation. Proc. Nat’l Acad. Sci. USA 115, E9550–E9559 (2018).

53. Alpert, A., Moore, L.S., Dubovik, T., Shen-Orr, S.S. Alignment of single-cell trajectories to compare cellular expression dynamics. Nature Meth. 15, 267–270 (2018).

54. Sanada, F., Taniyama, Y., Muratsu, J., Otsu, R., Shimizu, H., Rakugi, H., Morishita, R. IGF Binding protein 5 induces cell senescence. Front. Endocrinol. 9, 53 (2018).

55. Rocheteau, P. et al, A subpopulation of adult skeletal muscle stem cells retains all template DNA strands after cell division. Cell 148, 112–125 (2012).

56. Cornelison, D.D. and Wold, B.J. Single-cell analysis of regulatory gene expression in quiescent and activated mouse skeletal muscle satellite cells. Dev. Biol. 191, 270–283.

57. Tierney, M.T., Sacco, A. Satellite cell heterogeneity in skeletal muscle homeostasis. Trends Cell Biol. 26, 434–444 (2016).

58. Sanes, J.R., Lichtman, J.W. Development of the vertebrate neuromuscular junction. Annu. Rev. Neurosci. 22, 389–442 (1999).

59. Borisov, A.B., Dedkov, E.I., Carlson, B.M. Abortive myogenesis in denervated skeletal muscle: differentiative properties of satellite cells, their migration, and block of terminal differentiation. Anat Embryol (Berl) 209, 269–279 (2005).

60. Macpherson, P.C.D., Wang, X., Goldman, D. Myogenin regulates denervation-dependent muscle atrophy in mouse soleus muscle. J. Cell Biochem. 112, 2149–2159 (2011).

61. Borisov, A.B., Dedkov, E.I., Carlson, B.M. Differentiation of activated satellite cells in denervated muscle following single fusions in situ and in cell culture. Histochem Cell Biol 124, 13–23 (2005).

62. Xing, H., Zhou, M., Assinck, P., Liu, N. Electrical stimulation influences satellite cell differentiation after sciatic nerve crush injury in rats. Muscle & Nerve 51, 400–411 (2015).

63. Webster, M.T., Manor, U., Lippincott-Schwartz, J., Fan, C.M. Intravital imaging reveals ghost fibers as architectural units guidingin myogenic progenitors during regeneration. Cell Stem Cell 18, 243–252 (2016).

64. Stanganello, E., Hagemann, A.I.H., Mattes, B., Sinner, C., Meyen, D., Weber, S., Schug, A., Raz, E., Scholpp, S. Filopodia-based Wnt transport dueing vertebrate tissue patterning. Nature Commun. 6, 5846 (2015).

65. Goel, A.J., Rieder, M.K., Arnold, H.H., Radice, G.l., Krauss, R.S. Niche cadherins control the quiescence-to-activation transition in muscle stem cells. Cell Rep. 21, 2236–2250 (2017).

66. Prakash, S., Caldwell, J.C., Eberl, D.F., Clandinin, T.R. Drosophila N-cadherin mediates an attractive interaction between photoreceptor axons and their targets. Nature Neurosci. 8, 443–450 (2005).

67. Bonaldo, P., Sandri, M. Cellular and molecular mechanisms of muscle atrophy. Disease Models & Mech. 6, 25–39 (2013).

68. Bernet, J.D., Doles, J.D., Hall, J.K., Tanaka, K.K., Carter, T.A., Olwin, B.B. p38 MAPK signaling underlies a cell-autonomous loss of stem cell self-renewal in skeletal muscle of aged mice. Nature Med. 20, 265–271 (2014).

69. Budnick, V., Salinas, P.C. Wnt signaling during synaptic development and plasticity. Curr. Opin. Neurobiol. 21, 151–159 (2011).

70. Le Grand, F., Jones, A.E., Seale, V., Scime, A., Rudnicki, M.A. Wnt7a activates the planar cell polarity pathway to drive the symmetric expansion of satellite stem cells. Cell Stem Cell 4, 535–547 (2009).

71. Hall, A.C., Lucas, F.R., Salinas, P.C. Axonal remodeling and synaptic differentiation in the cerebellum is regulated by Wnt-7a signaling. Cell 100, 525–535 (2000).

72. Zhang, B. et al. Beta-catenin regulates acetylcholine receptor clustering in muscle cells through interaction with rapsyn. J. Neurosci. 27, 3968–3973 (2007).

73. Boers, H.E., Haroon, M., Le Grand, F., Bakker, A.D., Klein-Nulend, J., Jaspers, R.T. Mechanosensitivity of aged muscle stem cells. J. Orthop. Res. 36, 632–641 (2018).

74. Chen, F., Liu, Y., Suguira, Y., Allen, P.D., Greg, R.G., Lin, W. Neuromuscular synaptic patterning requires the function of skeletal muscle dihydropyridine receptors. Nature Neurosci. 14, 570–577 (2011).

75. Li, L. et al. Enzymatic activity of the scaffold protein Rapsyn for synapse formation. Neuron 92, 1007–1019 (2016).

76. Dell’Orso, S. et al. Single cell analysis of adult mouse skeletal muscle stem cells in homeostatic and regenerative conditions. Develop. 146, dev174177 (2019).

77. Giordani, L. et al. High-dimensional single-cell cartography reveals novel skeletal muscle-resident cell populations. Molec. Cell 74, 1–13 (2019).

78. Aguilar, C.A., Greising, S.M., Watts, A., Goldman, S.M., Peragallo, C., Zook, C., Larouche, J., Corona, B. Multiscale analysis of a regenerative therapy for treatment of volumetric muscle loss injury. Cell Death Discov. 4, 33 (2018).

